# Evolution of antigen-specific follicular helper T cell transcriptional programs across effector function and through to memory

**DOI:** 10.1101/2021.09.17.460841

**Authors:** Amanda M. Robinson, Brett W. Higgins, Andrew G. Shuparski, Karen B. Miller, Louise J. McHeyzer-Williams, Michael G. McHeyzer-Williams

## Abstract

Understanding how follicular helper T cells (TFH) regulate the specialization, maturation, and differentiation of adaptive B cell immunity is crucial for developing durable high-affinity immune protection. Using indexed-single cell molecular strategies, we reveal a skewed intra-clonal assortment of higher affinity TCR and the distinct molecular programming of the localized TFH compartment compared to emigrant conventional effector T_H_ (ETH) cells. We find a temporal shift in BCR class switch which permits identification of inflammatory and anti-inflammatory modules of transcriptional programming that subspecialize TFH function before and during the germinal center (GC) reaction. Late collapse of this local primary GC reaction reveals a persistent post-GC TFH population which discloses a putative memory TFH program. These studies define specialized antigen-specific TFH transcriptional programs that progressively direct class-specific evolution of high-affinity B cell immunity and uncover the transcriptional program of a memory TFH population as the regulators of antigen recall.

**Graphical Abstract:** 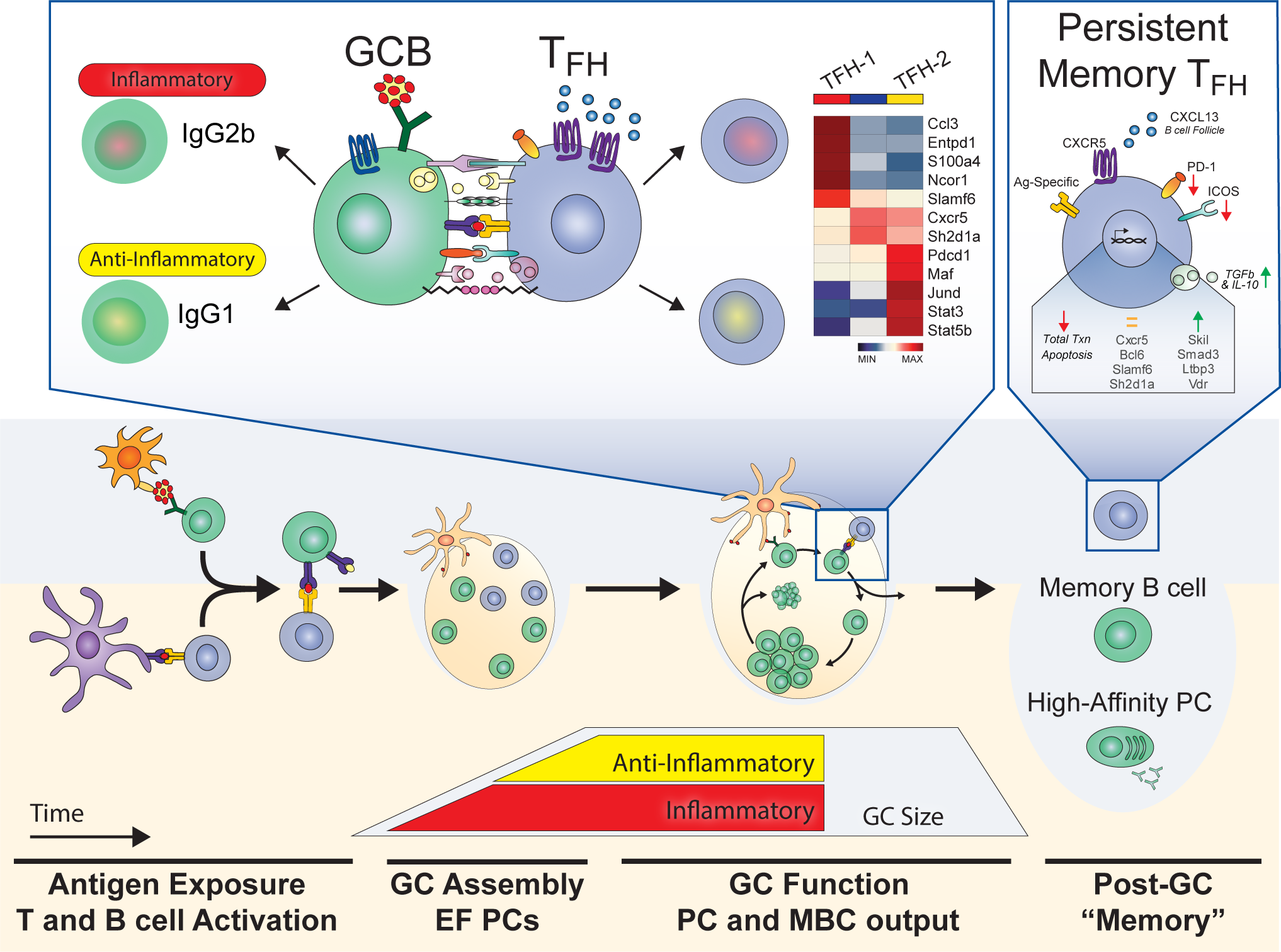

**Summary:** Distinct inflammatory and anti-inflammatory antigen-specific TFH transcriptional programs regulate class-specific B cell maturation.

**Highlights:**

- Skewed intra-clonal assortment of high affinity TCR into the TFH compartment
- Significant temporal delay in anti-inflammatory IgG1 production
- Inflammatory and anti-inflammatory transcriptional modules subspecialize TFH
- Late GC collapse reveals a persisting post-GC putative memory TFH compartment

## Introduction

Follicular helper T cells (TFH) are specialized to regulate the multi-faceted B cell response ultimately generating diverse antibody secreting plasma cells (PCs) and rapid recall ready memory B cells (MBCs) (*1*). Defined by their ability to migrate into the B cell follicle (*2*), TFH orchestrate the progressive developmental decisions of class-switch, germinal center (GC) affinity-maturation, and cell fate differentiation (*1, 3-7*). The class-switch event introduces functional heterogeneity: B cell receptor (BCR) classes offer separable effector functions which assort into conventional types of immunity: inflammatory type 1, anti-inflammatory type 2, and mucosal type 3 (*8*). Despite the distinct stepwise phases of B cell maturation and the established variability in the B cell response, TFH have been primarily studied as a single homogeneous population.

The TFH lineage is classically defined by expression of the chemokine receptor CXCR5 and the GC master transcription factor, Bcl6 (*9-11*). The duration and quality of antigen-specific cognate contact between TFH and B cells impacts the formation, function, and output from the GC (*4, 6, 12-15*). While B cells express unique class-specific transcriptional patterns (*16-18*), there is little understanding of how the cognate synapse differs and the TFH population is specialized to support the different B cell classes. Before TFH were discovered, conventional type 1 and type 2 helper T cell (TH_1_/TH_2_) associated cytokines, IL-4 and IFNγ respectively, were found to direct class-switch to inflammatory (IgG2b) and anti-inflammatory (IgG1) BCR subclasses (*19, 20*). While Bcl6 is a transcriptional repressor and inhibits TH_1_/TH_2_ development (*9-11*), TFH can express low levels of TH_1_/TH_2_ transcription factors, surface ligands, and cytokine signatures in specific exposure and autoimmune contexts (*21-28*). Furthermore, a subspecialized TFH population necessary for anaphylactic IgE production was recently discovered (*29*). IgE is a unique class of BCR in that it is the least abundant and primarily affiliated with immune dysfunction, namely allergy. Despite this evidence of an IgE-specific TFH subset, it remains unclear how TFH differentiate to direct the maturation of the predominant BCR classes in a healthy immune system.

The longevity of immune protection depends on the development of a robust antigen-specific memory compartment composed of PCs, MBCs, and memory TFH (*30*). Antigen-specific TFH are found in draining lymph nodes up to 200 days post immunization (*31*), but the TFH transition from primary effector function into persistent memory, as a transitional state or final memory fate, remains obscure. In humans and mice, circulating CXCR5+ CD4+ central memory T cells (TCM) can rapidly upregulate Bcl6, drive PC differentiation, and form secondary GCs (*21, 32-34*). However, the CXCR5+ TCM population can also give rise to secondary TH_1_ and TH_2_ populations in addition to TFH (*35*). In contrast to a pluripotent TCM reservoir, transgenic studies show TFH committed memory populations (*36*) and minimal lineage plasticity in splenic memory development (*37*). Notably, aged TFH were recently found to have heightened susceptibility to cell death which undoubtedly influenced previous studies (*38*). Understanding TFH transition from the primary GC response into the quiescent memory phase is crucial for the study of persistent immunological memory and the secondary response.

Here, we use indexed-single cell molecular strategies in a polyclonal antigen-specific model to resolve distinct stages of TFH function. Intraclonal assortment of transcriptionally distinct TFH and conventional effector CD4 T cells (ETH) demonstrated an affinity-based skewing of CD4+ T cell maturation. A temporal delay of type 2 anti-inflammatory IgG1 production revealed both an early TFH type 1 inflammatory pattern and a distinct, late arising, type 2 TFH transcriptional program. Lastly, significant collapse of the primary GC revealed a transcriptionally unique, putative local memory TFH population. Our studies provide insight into key stages of TFH function and uncover an array of targets for CD4 mediated class-specific and memory manipulation to inform future vaccine and immunotherapy design.

## Results

### Skewed intra-clonal assortment of PCC-specific TCR

To study a polyclonal CD4 T cell response, we use the well-established monovalent model protein antigen Pigeon Cytochrome C (PCC) (**Figure S1 A-C**). The T cell response to PCC is MHCII I-E^K^ restricted and readily identified by flow cytometry using the combination of Vα11 and Vβ3 T cell receptor (TCR) composition (*39*) and activation marker CD44 (**Figure 1 A**)(*31, 40*). As expected, we observed a significant expansion of PCC-specific T_H_ by day 7 that persisted through day 14 (**Figure 1 B**). Further classifying the PCC T_H_ response via L-selectin (CD62L) and CXCR5 expression, we subdivide conventional ETH (CD62L-, CXCR5-) and TFH compartments (CD62L-, CXCR5+); both followed the same pattern of local expansion after immunization (**Figure 1 A, B**). Furthermore, the TFH population expressed the highest level of PD-1 (**Figure 1 C**) and elevated levels of ICOS above the MPL background (**Figure S1 D**). By D14 the local TFH population had returned to naïve expression levels of CD43, an early activation marker that limits surface contact, while the ETH population maintained high CD43. The non-PCC “naïve” MPL background population was characterized as non-activated (CD44-, CD62L+) and non-Vα11Vβ3 though they were exposed to the TLR4 agonist adjuvant (monophosphoryl lipid A, MPL) via the immunization (**Figure S1 B**). This “naïve” MPL background population was also PD-1, ICOS, CD25 and CD43 deficient, while positive for the naïve and memory T cell marker, Ly6c (*41*) (**Figure S1 B, D**).

**Fig 1.**
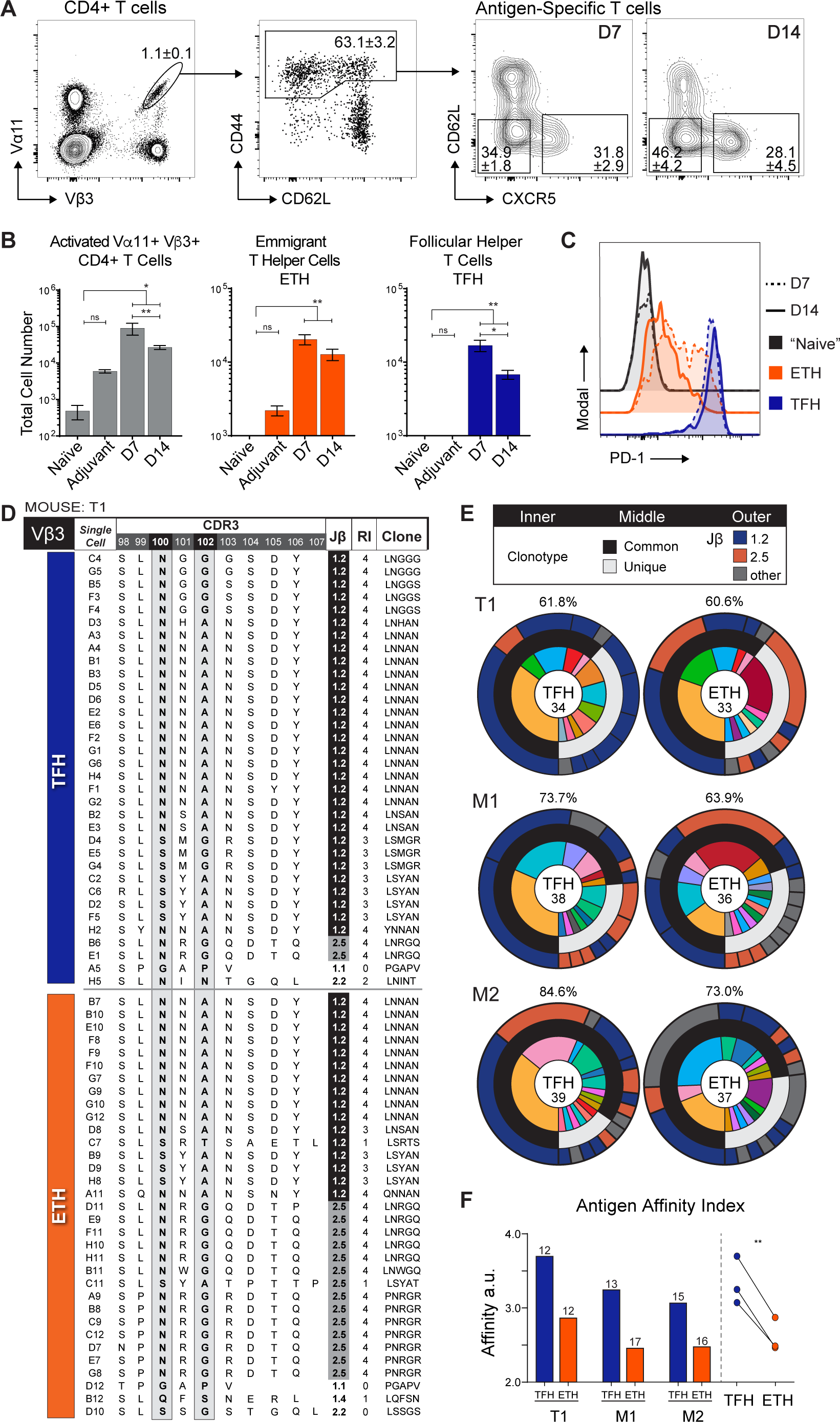
Skewed intra-clonal assortment of PCC-specific TCR. **(A)** Flow cytometry in draining LNs 7 days after base of tail immunization with NP-PCC in MPL. First pannel is CD4 T cells (CD4+, B220-, CD8-), second pannel is Va11+ Vb3+ CD4 T cells, third pannels are PCC specific T cells (CD4+, Va11+ Vb3+ CD44+). D7 and D14 as indicated. Numbers in outlined areas indicate frequency of gated population (mean ± s.e.m.). **(B)** Quantification of antigen-specific T cells, conventional effector T helper cells (ETH; CD62L-, CXCR5-), and follicular helper T cells (TFH; CD62L-, CXCR5+). Statistics two-tailed Mann-Whitney t.test. * P≤0.05, ** P≤0.01. **(C)** Histogram expression of PD-1 by non-specific “naïve” (Va11-, Vb3-, CD62L+, CD44-), ETH, and TFH. **(D)** Vβ3 CDR3 region sequences of index sorted Vα11 Vβ3 CD44hi, CD62lo, TFH (CXCR5+) and ETH (CXCR5-) from draining LNs 7 days after base of tail immunization with PCC in MPL. Jβ usage indicated; high affinity 1.2, low affinity 2.5. Repertoire index (RI) is the sum of canonical PCC specific features. **(E)** Clonal composition of indicated population (inner ring), commonality of clones (middle ring), and Jβ usage (outer ring). Phenotype and number of cells sequenced listed in center. Percentage listed above is the percentage of each population composed of common clones. Paired TFH and ETH populations from individual mice displayed side by side. **(F)** Population affinity index of each population, number listed is total number of clonotypes. log2([(#Jb 1.2 *4)+(#Jb 2.5*2)+(#Jb other)]/total). Statistics two-tailed ratio paired t.test. * P≤0.05, ** P≤0.01. Data are representative of four to ten experiments (A-C; mean and s.e.m. of n=4-16 per timepoint) or one experiment of three mice (D-F).

To investigate clonal dispersion among PCC-responding TH, ETH and TFH were index sorted and the Vβ3 CDR3 junction was sequenced (**Figure 1 D**, **S1 E**). Four PCC-specific specific β-chain features have been extensively characterized: asparagine at position 100, alanine or glycine at position 102, 9 amino acid CDR3, and Jβ chain 1.2 (high affinity) or 2.5 (low affinity) (*40, 42*). Attesting to the accuracy of TCR based identification, 94% of sorted Vα11Vβ3 TFH exhibited a minimum of three PCC canonical features. ETH and TFH populations exhibited notable (30-40%) clonal similarity (**Figure 1 E**, **inner ring**) and these shared clonotypes expanded to constitute 60-85% of the sequenced ETH and TFH populations (**Figure 1 E**, **middle ring**). This suggests naïve CD4 clones are receptive to external stimuli and not intrinsically predestined for specific maturation fates.

We previously found one such influential stimuli is TCR affinity for antigen via tetramer binding (*43*). Using the established difference in affinity due to Jβ usage (*42*), we uncovered a distinct intraclonal skewing of affinity toward the TFH fate. While clones using Jβ 1.2 and 2.5 were common in both populations (**Figure 1 E**, **outer ring**), the combinatorial population affinity index was consistently higher in TFH populations relative to their ETH counterparts (**Figure 1 F**). Notably, this high-affinity skewing did not diminish the clonal diversity of the TFH. The paired TFH and ETH populations contained a nearly equivalent number of unique clones despite the noted difference in affinity (**Figure 1 F**, unique clone # listed above bars). Overall, these data establish the production of a clonally diverse high-affinity TFH population driving a B cell response to PCC immunization.

### Inflammatory programming predominant in the early TFH compartment

To investigate the specific transcriptional patterns establishing specialized TFH functionality, we utilized a custom quantitative and targeted single-cell sequencing protocol to achieve higher sensitivity for selected mRNA species (*44, 45*). First, we index sorted day 7 antigen-specific ETH and TFH along with “naive” (non-specific MPL background CD44lo CD62Lhi) T cells (**Figure 2 A**). As expected, the index sort confirmed the TFH population exhibited the highest levels of PD-1 (**Figure 2 B**). Individual cells from these sorted compartments were uniquely bar-coded during cDNA synthesis prior to a targeted and nested PCR amplification of ∼500 mRNA species before digitally deconvoluting the multiplexed data for single cell quantification (qtSEQ) (**Figure S2 A-E**). We see an average UMI coverage of 330 amplified reads per original UMI, with technical thresholds set at a minimum of 1000 reads, a minimum of 60 separate UMIs and a maximum of 2000 UMIs per cell (**Figure S2 B-D**).

**Fig 2.**
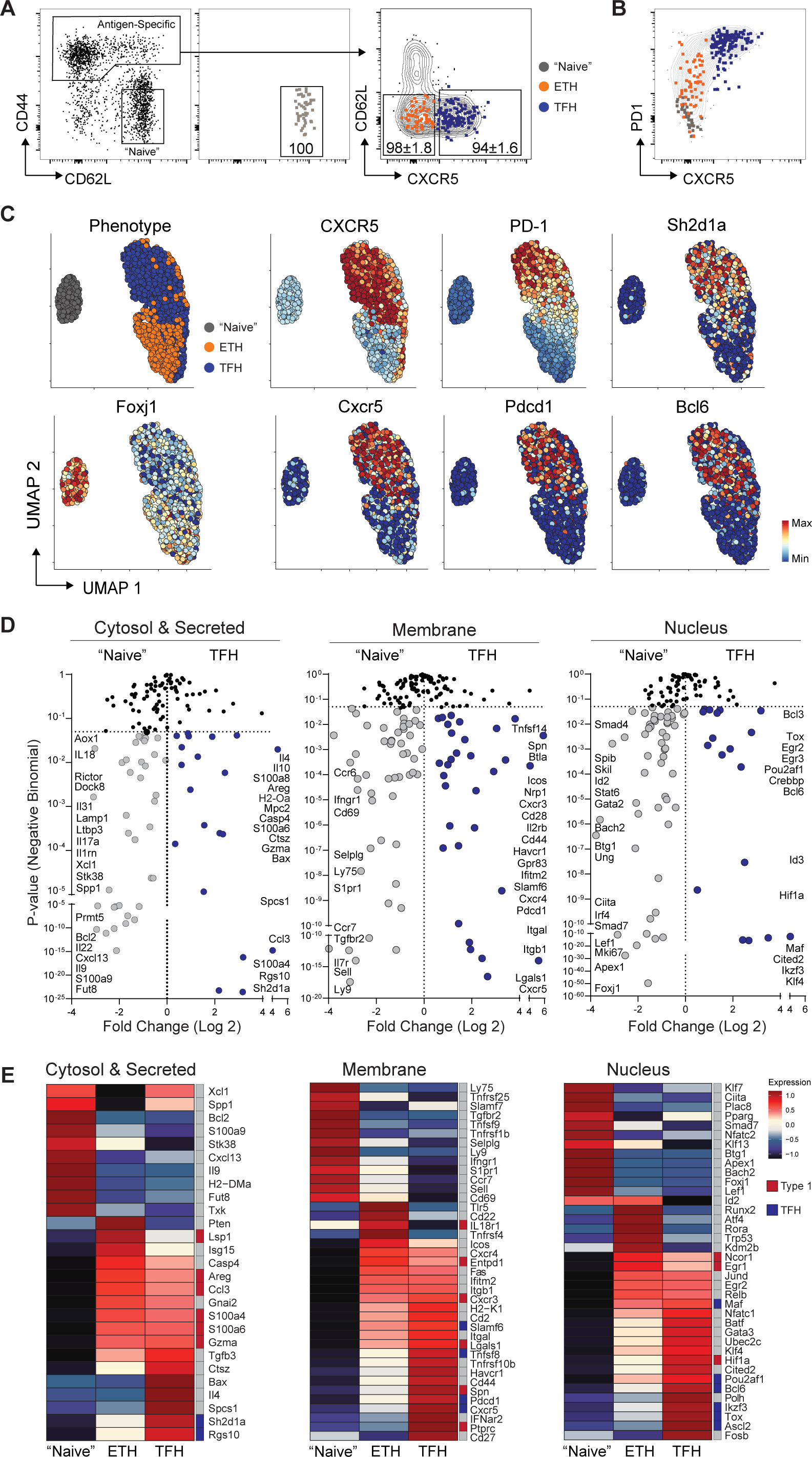
Inflammatory programming predominant in the early TFH compartment. **(A)** Representative index sorted populations from draining LNs 7 days after base of tail immunization with NP-PCC in MPL. Grey, “Naïve” (CD62L+, CD44-). Orange, ETH (CD44+, CD62L-, CXCR5-). Blue, TFH (CD44+, CD62L-, CXCR5+). Numbers in outlined areas indicate frequency of sorted population in the gates (mean ± s.e.m.). **(B)** Representative PD-1 and CXCR5 expression of index sorted populations. Contours represent total PCC specific CD4 T cells (Va11+ Vb3+ CD44+). **(C)** UMAP dimensionality reduction of index sorted cells. Surface protein and transcript expression overlayed as labeled. **(D)** Differential gene expression between “naïve” and TFH populations separated by designated location of function. Fold change plotted against negative binomial P.value. **(E)** Pseudobulk gene expression of top 20 variable genes of each population, total 57 genes. Genes assorted by functional location, color scale determined by z score distribution, literature associated expression and/or effector function indicated (Type1, red; TFH, blue). Data are representative of three separate experiments, n=12. Sorted cells; “Naïve” n= 214, ETH n=191, TFH n=318.

UMAP dimensionality reduction of the indexed surface proteins demonstrated a clustering of antigen-specific T cells apart from the naïve population, as well as a regional distinction between ETH and TFH (**Figure 2 C**). Overlaying the protein and mRNA expression of conventional TFH identifiers on the UMAP coordinates revealed a consistency in protein and mRNA expression patterns across CXCR5, PD-1 (**Figure 2 C**), and ICOS (**Figure S2 F**). Additionally, TFH associated *Bcl6, Sh2d1a* (SLAM), and *Rgs10* were prevalent among TFH, while *Foxj1* was most prevalent in “naïve” and, to a lesser extent, ETH regions (**Figure 2 C**, **S2 F**). It is of note that this “naïve” population was collected following the naïve T cell gating strategy (**Figure S1 B**) from immunized mice and thus controls for the effects of innate TLR4 stimulation via the adjuvant MPL (**Figure S2 G**). While this adjuvant background stimulation is robust, we see antigen-specific response indicators (upregulated *CD44, ICOS*; downregulation *Ccr7, Sell, Foxj1*) consistent in the antigen-specific ETH and TFH compartments (**Figure 2 D, E**, **S2 H**).

The capacity for follicular localization (*Cxcr5*) and increased surface engagement (*Cd27, Cd28, Cd40lg, Lgals1, Itgal, ICOS, Nrp1, Slamf6*) distinguish the early day 7 TFH (**Figure 2 A, D, E**, **S2 H**). Furthermore, the upregulation of transcription factors (*Pou2af1* (Bob1)*, Tox, Ascl2, and Ikzf3* (Aiolos)) indicate an early commitment to the upregulation of Bcl6 and the TFH fate. Interestingly, in addition to the co-inhibitory receptor *Pdcd1*, early TFH have upregulated other inhibitory receptors (*Btla*, *Havcr1*) and negative growth regulator (*Lgals1*) potentially contributing to tight control of the TFH compartment.

Despite the robust follicular program, further scrutiny of the targeted transcriptional patterns revealed a shared inflammatory antigen-specific program among ETH and TFH. We see elevated levels of cytosolic genes (*Ccl3, Lsp1, Gzma, Areg),* secreted S100 proteins (*S100a4, S100a6, S100a8*), T_H_1 associated surface receptors (*Cxcr3*, *Ptprc* (CD45)*, Entpd1* (CD39), *Lgals1*) and antigen-specific membrane bound activation marker genes (*Cd44, Spn*) (**Figure 2 D, E**). The shared upregulation of type 1 associated program is reflective of the inflammatory environment exacerbated by the adjuvant (MPL) and we propose these inflammatory features are directing downstream inflammatory B cell class-switching and maturation.

### Distinct temporal delay in anti-inflammatory IgG1 B cell response

While PCC is an advantageous model when studying CD4 T cells, the PCC-specific B cell response is elusive. High-affinity PCC-specific B cell responses may by prevented through processes evolved to avoid autoimmunity – cytochrome C being a highly conserved component of the mitochondrial electron transport chain. To circumvent this challenge, we conjugated the well-studied small hapten molecule, 4-hydroxy-3-nitrophenylacetyl (NP) to PCC as the carrier protein. Following subcutaneous immunization with NP-PCC in MPL adjuvant, we find a substantial antigen-specific class-switched B cell response is present by day 7 and continues through day 14 (**Figure 3 A, B**). The B cell response can be further characterized as antigen specific plasma cells (PCs; CD138+), germinal center B cells (GCBs; B220+, GL7+, CD38-), and memory B cells (MBCs; B220+, GL7-, CD38+) (**Figure 3 C-F**, **S3 A**). The day 7 B cell response is dominated by PC production, while the day 14 response demonstrates a robust GC response producing both PCs and MBCs (**Figure 3 G**).

**Fig 3.**
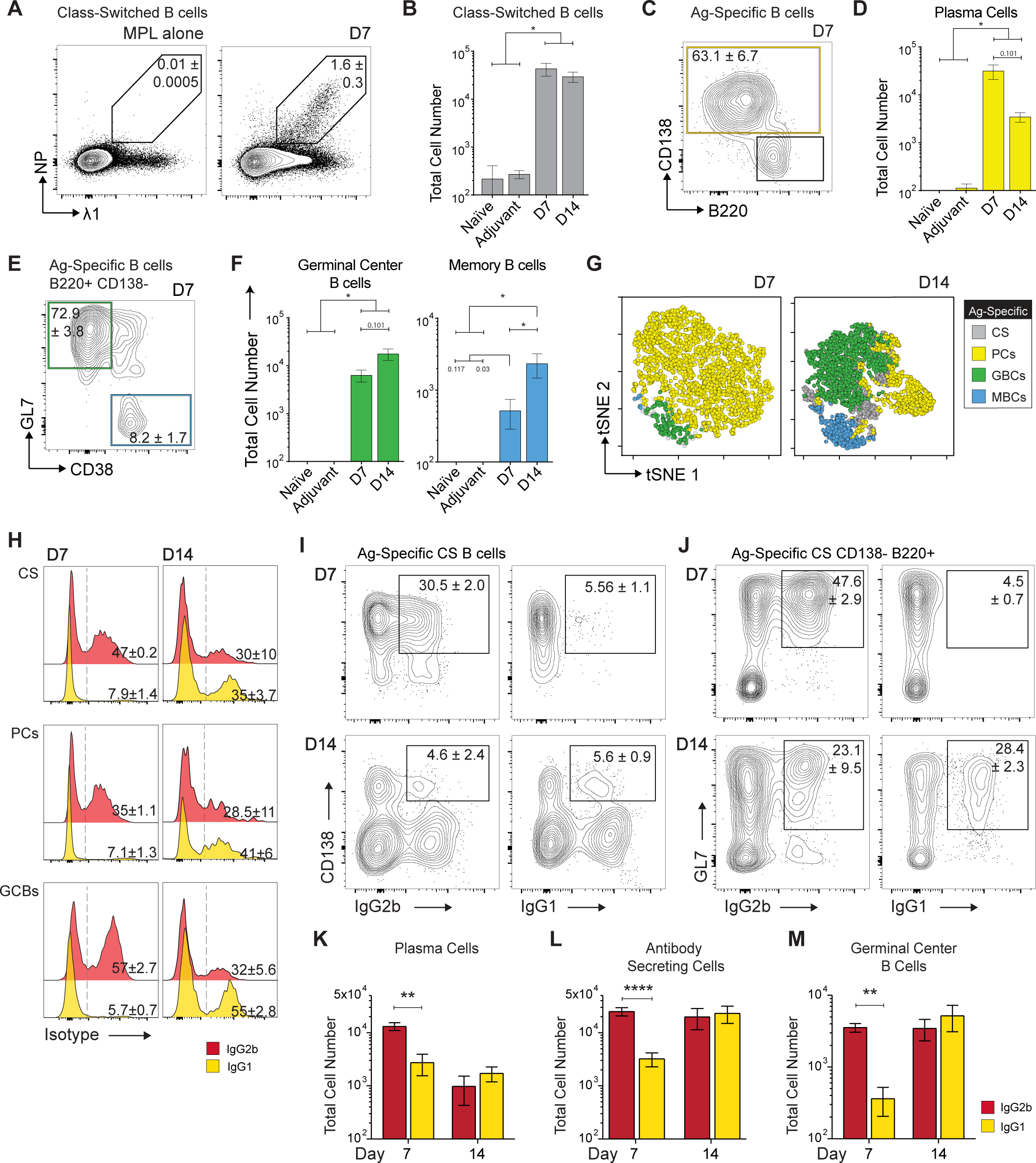
Distinct temporal delay in anti-inflammatory IgG1 B cell response. **(A)** NP specific B cell gating strategy in draining lymph nodes following NP-PCC immunization. Representative flow plot from day 7, concatenated file of 3 mice. **(B)** Total cell number of antigen-specific B cells. **(C)** NP specific plasma cell gating strategy. **(D)** Total cell number of antigen-specific PCs. **(E)** Gating strategy for antigen-specific germinal center B cells (GCBs) and memory B cells (MCBs). **(F)** Total cell number of Ag-specific GCBs and MBCs. **(G)** tSNE dimensionality reduction of CS Ag-specific B cell response at day 7 and day 14. Colors indicate phenotype; grey, Class switched B cells (CS); yellow, Plasma Cells (PCs); green, Germinal Center B Cells (GCBs); blue, Memory B cells (MBCs). **(H)** Concatenated modal histogram of isotype expression at day 7 and day 14 (yellow, IgG1; red, IgG2b) among CS NP+ B cells, PCs, and GCBs. **(I)** FACS expression of CD138 and IgG1 or IgG2b of CS NP+ B cells. **(J)** FACS expression of GL7 and IgG1 or IgG2b of CS NP+ CD138-B220+ B cells. Representative flow plots (I, J), concatenated files of 3 mice. **(K)** Total cell number of IgG1 and IgG2b PCs. **(L)** Bulk elispot for NP-specific antibody secreting cells (ASC). **(M)** Total cell number of IgG1 and IgG2b GCBs. Statistics two-tailed Mann-Whitney t.test. * P≤0.05, ** P≤0.01. Data are representative of four to seven experiments (mean and s.e.m. of n=3-10 per timepoint).

Having established concurrent antigen-specific B and T cell access, we interrogated the PCC driven B cell response with isotype specificity to uncover the impact of this type 1 presenting day 7 TFH population. The day 7 early class-switched B cell response was dominated by inflammatory IgG2b B cell products relative to anti-inflammatory IgG1 (**Figure 3 H**). The IgG2b response was ten-fold larger than the IgG1 response at day 7 (**Figure 3 H**). The predominant IgG2b response ranged from 35-60% of the total class-switched B cells, PCs, and GCBs at day 7 (**Figure 3 H-M**, **S3 B**). This early inflammatory predominance persisted slightly longer than the observed waves of IgG2b and IgG1 production seen in the response to NP-KLH with MPL (*18*). However, by day 14, the anti-inflammatory IgG1 population had expanded to be equivalent to, if not greater than, the inflammatory IgG2b population in both frequency and total cell numbers (**Figure 3 H-M**, **S3 B**). Curiously, the production of IgG1 at day 14 is not uniform across the draining lymph node, but rather regionally expanded (**Figure S3 C-H**). Taken together, the early predominance of IgG2b mirroring the inflammatory program in the day 7 TFH and the delayed, topographically skewed, IgG1 expansion indicate these waves of functionally specific B cell fates require differential TFH support for differentiation and maturation.

### Inflammatory and anti-inflammatory transcriptional patterns subspecialize TFH

We used this delay in anti-inflammatory B cell activity to explore the TFH population for transcriptional programs specifically directing the early type 1 IgG2b and late arising type 2 IgG1 responses. UMAP analysis of naïve T cells, day 7 TFH, and day 14 TFH revealed a separation of antigen-specific TFH apart from the non-specific naïve population; however the day 7 and day 14 TFH were mixed across the TFH region (**Figure 4 A**). Furthermore, the UMAP analysis generated 5 transcriptionally distinct clusters, three of which composed the majority of the TFH UMAP region (**Figure 4 B**, **S4 A**).

**Fig 4.**
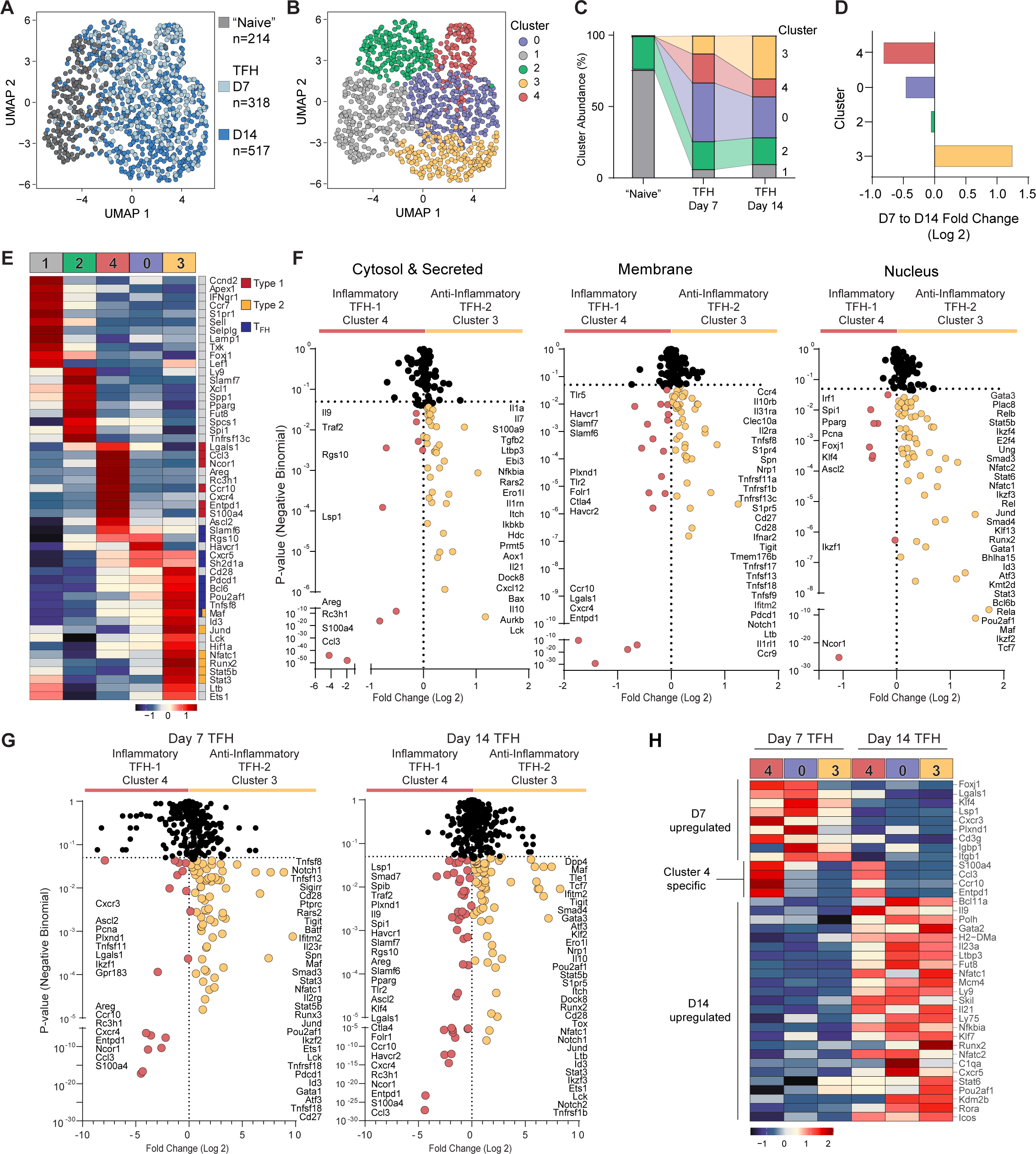
Inflammatory and anti-inflammatory transcriptional patterns subspecialize TFH. **(A)** UMAP dimensionality reduction of “naïve” (grey, n=214), day 7 PCC-specific TFH (light blue, n=318), day 14 PCC-specific TFH (blue, n=517). **(B)** UMAP generated clusters (Euclidean Distance). **(C)** Proportion of each timepoint sample allocated to UMAP clusters (cluster abundance). **(D)** Fold change in the abundance of each cluster at day 7 or day 14. (Log 2(D14/D7)). **(E)** Pseudobulk Gene Expression Heatmap; Negative Binomial Fold >0.25, P <0.05 in a minimum of 3 differential gene comparisons and among top 20 variable genes. Literature associated expression and/or effector function indicated (Type1, red; Type 2, yellow; TFH, blue). **(F)** Differential gene expression of cluster 3 and cluster 4. Genes divided across functional locations. **(G)** Differential expression of cluster 3 and cluster 4 at day 7 and day 14. **(H)** Pseudobulk heatmap of genes differentially expressed at day 7 and day 14 across three TFH clusters. Data are representative of three separate experiments, n=12. Sorted cells; “Naïve” n= 214, D7 TFH n=318, D14 TFH n=517.

The abundance of each cluster at the sorted timepoints revealed the entirety of the naïve sorted population was represented by cluster 1 and 2, while the majority of the day 7 and day 14 TFH populations were split among clusters 0, 3 and 4 (**Figure 4 C**). As expected, cluster 1 (grey) and cluster 2 (green), composing the naïve population, have the highest expression of naive T cell markers (*Ccr7, Sell, Foxj1, Lef1*). Interestingly, cluster 2, and to a lesser extent cluster 1, express a set of genes increasing surface contact (*Ly9, Slamf7*), T cell activation mediators (*Fut8, Pparg*) and protein synthesis machinery (*Spcs1*) suggesting, these are MPL agonized naïve T cells or possibly contain a quiescent background memory subset. Nevertheless, both cluster 1 and 2 lack expression of the classic antigen-activated T cell or TFH markers (**Figure 4E, S4 C, D**).

Genes historically associated with TFH function (*Cxcr5, Bcl6, Pdcd1, Slamf6, Icos, Sh2d1a*) are expressed by all three TFH clusters (cluster 4-red, cluster 0-blue, cluster 3-yellow) (**Figure 4 E**, **S4 C, D**). Cluster 0 (blue) appears to represent a baseline TFH population without additional cluster unique transcription (**Figure 4 E**, **S4 C**). Determined by the fold change of each cluster from day 7 to day 14, this baseline TFH exhibited the smallest shift in prevalence, while clusters 3 and 4 were notably skewed toward day 7 or day 14 TFH (**Figure 4 D**). Cluster 4 (red) is nearly twice as prevalent in the day 7 TFH and we hypothesize this cluster is the subspecialized inflammatory TFH as we saw a predominance of inflammatory B cell products at day 7. Conversely, only cluster 3 (yellow) expanded from day 7 to day 14, more than doubling in frequency. As the only cluster to expand, we hypothesize this population is responsible for supporting the delayed anti-inflammatory IgG1 response. Further supporting a relationship between the IgG1 response and cluster 3 TFH, the frequency of cluster 3 (yellow) maintains a similar proportion to the frequency of IgG1 products at day 7 and day 14 while cluster 0 (blue) and 4 (red) exhibit wide variability (**Figure S4 B**).

In the day 7 predominant cluster 4 (red), there is significant shared upregulation of type 1 immunity associated genes (*Ccl3, S100a4, Ccr10, Entpd1, Lsp1, Ncor1, Lgals1*) consistent with the early day 7 response (**Figure 4 F, G**). Furthermore, additional T_H_1 and inflammation linked genes (*Tlr5, Tlr2, Plxnd1*) complement the shared core TFH program. The upregulation of inhibitory co-receptors (*Havcr1, Havcr2, Ctla4*) potentially contribute to a dampening of inflammatory B cell production as we see the frequency of IgG2b products declining. However, because the day 7 timepoint is simultaneously dominated by IgG2b production and PC differentiation, we looked at the differential gene expression across the TFH within each timepoint to determine which features were contributing to inflammatory transcriptional programming or more likely directing the PC fate (**Figure 4 H**). A number of inflammation-associated genes (*S100a4, Ccl3, Ccr10, Entpd1*) are limited to our hypothesized inflammatory type 1 cluster 4 (**Figure 4 H**). However, early day 7 TFH had consistently higher expression of some inflammatory genes (*Lsp1, Cxcr3, Lgals1*), in addition to surface molecules (*Plxnd1, Igtb1*) and transcription factors (*Foxj1, Klf4*). Thus, because the clusters unaffiliated with the inflammatory response (cluster 0, blue; cluster 3, yellow) similarly upregulate these genes at day 7, they are likely contributing to PC differentiation rather than inflammatory function. Similarly, a breadth of genes are consistently expressed at higher levels in day 14 TFH; we see higher expression of *Cxcr5*, *Il-21*, and *Icos* – indicative of GC location and function – along with potentially novel GC markers (*Bcl11a, Nfatc1, Nfact2, Mcm4, Ly9, Ly75, Klf7*) (**Figure 4 H**).

We next probed the cluster 3, which expanded from day 7 to day 14 with the anti-inflammatory IgG1 response and uncovered a systematic type 2 program. TH_2_ affiliated signaling modulators (*Dock8, Ebi3*), secreted factors (*Il-10, Tgfb2*), cell surface ligands (*Ifitm2, Nrp1, Tigit, S1pr4, S1pr5*), and transcription factors (*Gata3, Jund, Stat3, Stat5b, Maf, Nfatc1, Runx2*) are upregulated in cluster 3 (**Figure 4 E-G**, **S4 D**). Notably, *Gata3*, the master transcription factor of the TH_2_ lineage, and *Dock8*, the key knock-out gene underlying the identification of the IgE regulating TFH-13 subset are among those upregulated (*29*). Furthermore, many TNF family members (*Cd27, Ltb, Tnfsf8, Tnfsf9, Tnfsf18, Tnfsf13, Tnfrsf17, Tnfrsf13c, Tnfrsf1b, Tnfrsf11a*), signaling mediators (*Lck, Rc3h1*) and additional cytokines (*Cxcl12, Il-21, Il7, Il1a*) are increased in cluster 3. The expression of *Il-21* and other TFH associated genes (*Pdcd1, Pou2af1, Tnfsf8*) demonstrates the complementary expression of follicular and anti-inflammatory transcriptional programs. Taken together, these diverging type 1 and type 2 transcriptional programs supplementing a core follicular program indicate that the TFH population is subspecialized to orchestrate the B cell response with BCR class specificity.

### TFH population survives GC collapse and segregates from conventional CD4+ Memory

The primary antigen-specific GC collapses by day 28 with only 10% of the day 14 GC population remaining (**Figure 5 A, B**). Despite this local GC collapse, the antigen-specific TFH population remained, only contracting to half the original size (**Figure 5 C, D**). Previous work in the PCC system established that the majority of antigen-specific follicular T cells at day 7 are not within a GC; however, by day 9, when the GC is established and productive, the follicular T cells are predominantly in a GC (*42*). We found antigen-specific TFH outnumber GCBs at day 7, the primarily non-GC timepoint, while this ratio flipped by GC dominant timepoints, day 10 and day 14 (**Figure 5 E**). At the late day 28 timepoint, antigen-specific TFH again outnumber GCBs as seen at the day 7 follicular non-GC timepoint (**Figure 5 E**). This swing in relative population size suggests the majority of these TFH are again residing in the follicle but not in a GC and therefore no longer supporting GC function.

**Fig 5.**
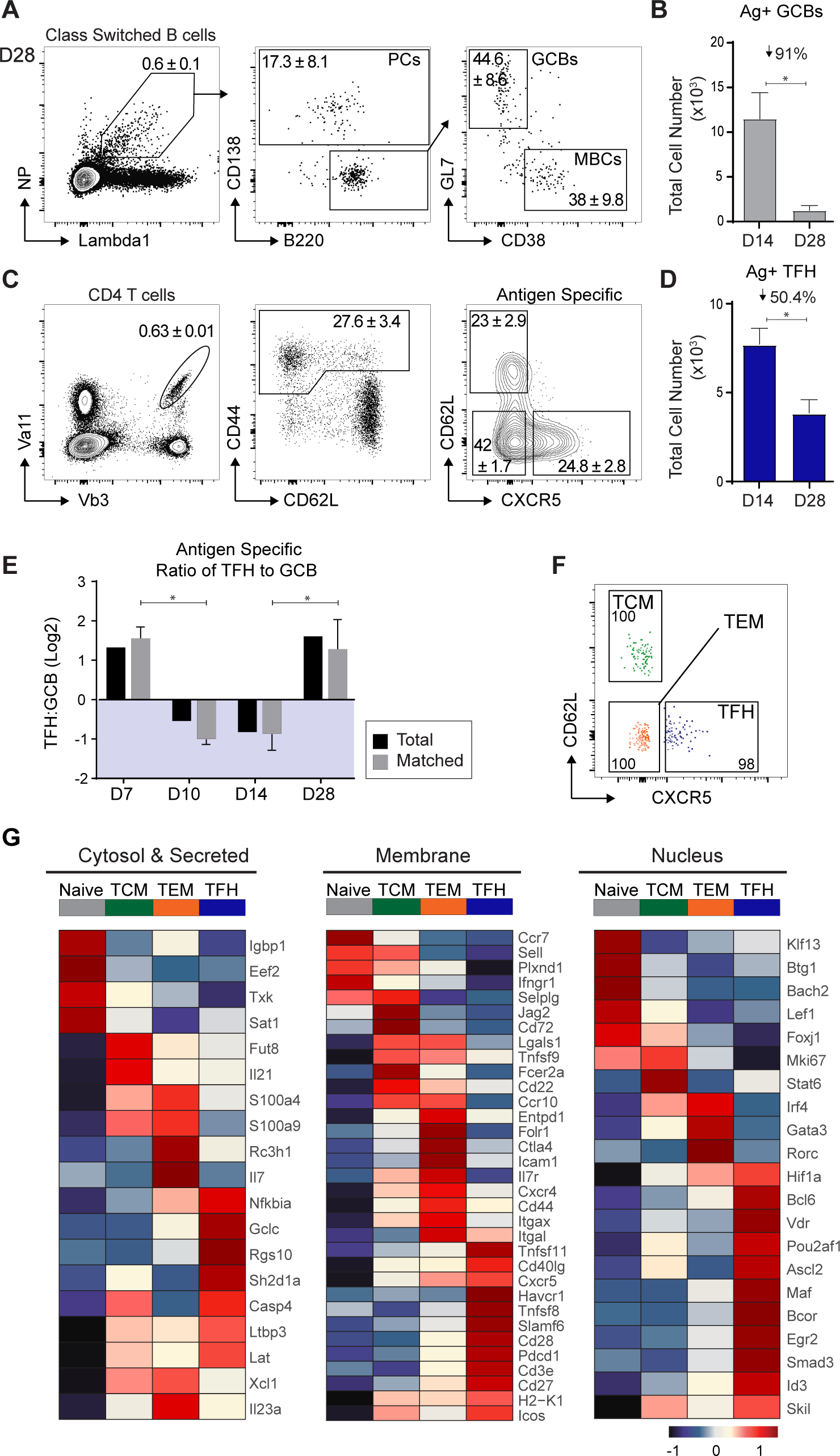
TFH population survives GC collapse and segregates from conventional CD4+. **Memory (A, B)** Flow identification (A) and total cell number (B) of antigen-specific GCBs at D28. Percentage indicated is size of GCB population loss. **(C, D)** Flow identification (C) and total cell number (D) of TFH at D28. Percentage indicated is size of TFH population loss. **(E)** Ratio of Ag-specific TFH:GCBs throughout the NP-PCC response. Matched indicates TFH and GCBs from same mice (7 n=2, 10 n=3, 14 n=3, 28 n=3). Two Tailed Mann-Whitney t.test of matched data. **(F)** Index sort Central Memory (TCM), Effector Memory (TEM), and putative TFH memory (TFH). **(G)** Pseudobulk heatmap expression of top 30 variable genes of each population across “naïve”, TCM, TEM, and TFH populations. Heatmaps split by functional location of the gene products.

Next, we investigated this putative post-GC memory TFH within the context of conventionally studied CD4 central (TCM) and effector (TEM) memory populations (**Figure 5 F**). The post-GC TFH maintains the highest levels of PD-1 expression but shares low expression of surface ligand ICOS and anti-adhesion molecule CD43 with TEM and TCM memory populations (**Figure S5 A**). We also see levels of Ly6C trending towards naïve levels, while entirely absent in the primary TFH and ETH precursor populations (**Figure S5 A**).

Deepening our analysis, we assessed similarities and differences in the memory populations against the naïve and MPL background control T cell groups (**Figure S5 B**). Cytosolic and secreted molecules (*S100a4, S100a9, Ccl3*, *Rch31)* and surface ligands (*Ltb, Lgals1, Cd44, Cxcr3, Ccr10*) formerly present in primary TFH are now predominantly expressed by day 28 TCM and TEM compartments (**Figure 5 G**, **S5 C**) suggesting the follicular environment is no longer inflammatory. However, primary TFH markers (*Cxcr5, Pdcd1, Slamf6, Sh2d1a, CD40lg, Pou2af1, Bcl6*) are upregulated in the local day 28 TFH population maintaining B cell proximity and capacity for surface engagement (**Figure 5 G**, **S5 C, D**). Further distinguishing the putative memory TFH are inhibitory molecules (*Rgs10, Tnfsf8, Havcr1, Sigirr* (TIR8)), signaling modifiers (*Lat, Id3, Bcor*) and genes implicated in metabolic and cell cycle regulation (*Gclc, Rgs10, Casp4, CD24, CD27, Tnfsf11*) (**Figure 5 G**, **S5 C, D**). The observed transcriptional restriction of genes with inhibitory function in day 28 TFH reflect the recent shutting down of active effector function and collapse of the functional GC. In addition to the upregulation of genes limiting cellular proliferation and survival, the lowest levels of *Mki67* expression (**Figure 5 G**, **S5 C**) indicate day 28 TFH have recently and are not currently proliferating. Taken together, we consider this distinct post-GC transcriptional program indicative of a shift in TFH function concluding the primary GC response.

### Transition to memory TFH initiates PD-1 downregulation

With respect to the primary day 7 and day 14 TFH, we see conserved yet lower expression of primary TFH markers (*Cxcr5, Pdcd1, Icos, Cd44, Pou2af1, Bcl6, Slamf6, Sh2d1a, Rgs10)* in day 28 TFH (**Figure 6 A, B**, **S6 A**). The maintained downregulation of *Ccr7* and *Sell* suggest the day 28 TFH are remaining in the B cell follicle, not returning to the T cell zone nor recirculating (**Figure 6 A, B**). Distinguishing the persistent TFH apart from primary TFH are TNF family members (*Tnfsf11, Tnfsf8, Tnfsf9, Cd27*), signaling modifiers (*Smad3, Ltbp3)*, interleukins (*Il-2, Il-10, Il-4*), and transcription factors (*Skil, Vdr)* (**Figure 6 A, B**). Interestingly, the co-stimulatory ligand *Cd28* is most highly expressed among post-GC TFH, while the expression of the direct competitive-inhibitor, *Ctla4*, is maintained from day 14 to day 28. This implies a reduction of *Ctla4* mediated inhibition, thus effectively lowering the signaling threshold for post-GC TFH. Furthermore, this day 28 transcriptional profile is separable from the UMAP clusters subspecializing the day 7 and day 14 TFH (**Figure S6 B**). Notably, the day 28 TFH cluster is most similar to Cluster 0 (blue) which previously appeared to represent the baseline TFH population suggesting the transition into memory returns to and maintains a core follicular program (**Figure S6 C**). As the persistent TFH express a separable transcriptional program after the collapse of GC we propose this program is indicative of the transition out of primary TFH function.

**Fig 6.**
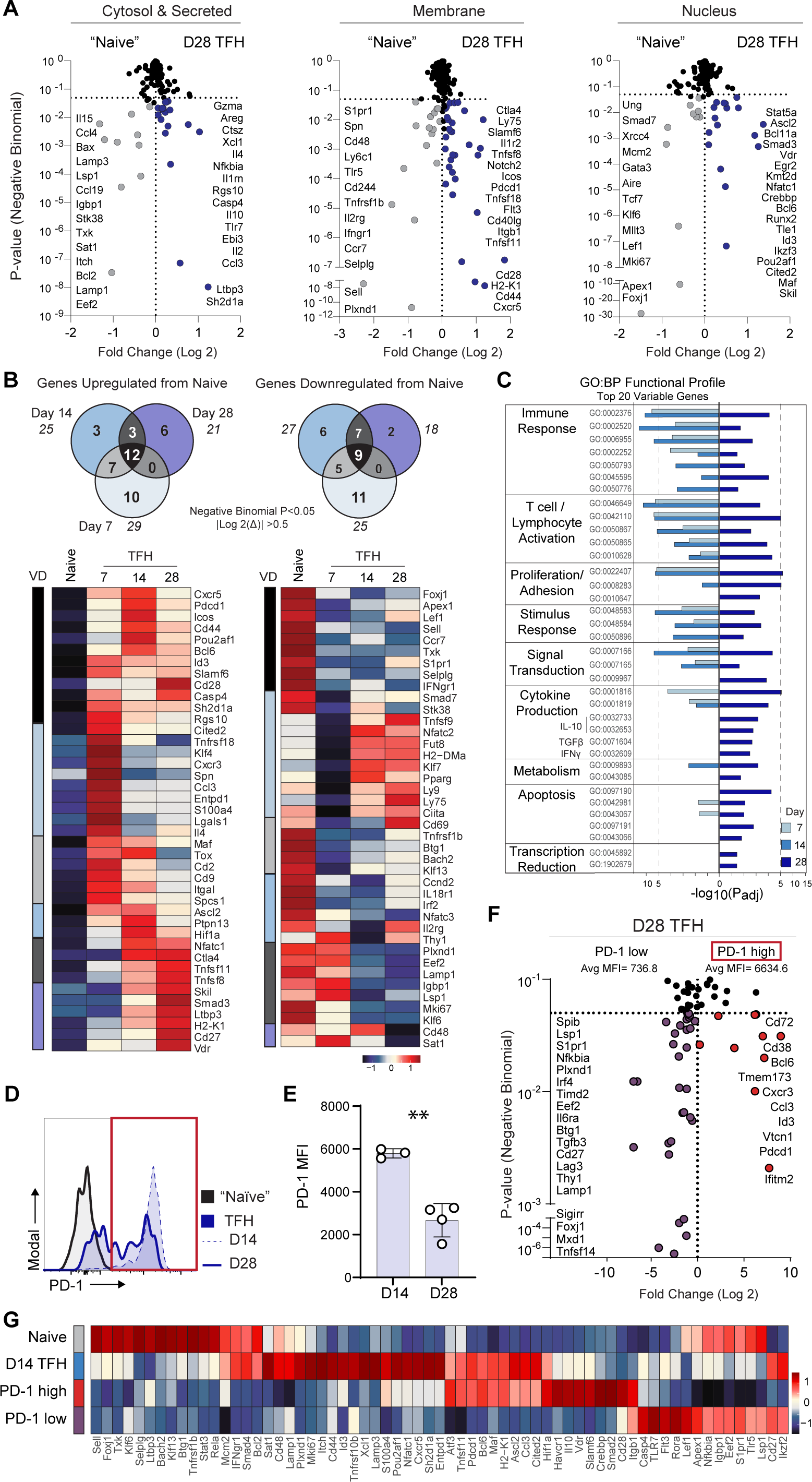
Transition to memory TFH initiates PD-1 downregulation. **(A)** Differential gene expression between MPL alone and D28 TFH populations separated by designated location/function. Fold change plotted against negative binomial P-value. **(B)** Venn Diagram of shared and differentially expressed genes across D7, D14, and D28 TFH with respect to the Naive CD44lo CD62Lhi population. Genes have P-value <0.05 and fold change >0.5. Italic number listed is the sum genes up or down regulated by that population. Below, pseudobulk heatmap of genes listed in up/down-regulated venn-diagram. VD column corresponds to the area of the venn-diagram represented. Heatmap of Naive, D7, D14, and D28 TFH. **(C)** gProfiler gene enrichment of GO Biological Process (GO:BP) pathways. Input genes were top 20 variable genes of day 7, D14, and D28 TFH populations. P-value is indicated by distance from 0; right/left is formatted for clarity. **(D)** Representative histogram of PD-1 surface expression. D14 n=6, D28 n=7. **(E)** Geometric mean PD-1 expression at D14 and D28. D14 n=3, D28 n=4. Statistics, Two-tailed Mann-Whitney t.test, * P≤0.05, ** P≤0.01. **(F)** Differential gene expression of PD-1 high vs PD-1 low D28 TFH. (PD-1 high MFI>2000). **(G)** Pseudobulk heatmap of top variably expressed genes across PD-1 high and low populations

Biological process identification via GO enrichment analysis of the top 20 variably expressed genes at day 7, day 14, and day 28 revealed shared follicular functions and unique post-GC features (**Figure 6 C**). First, a common set of biological processes associated with the general immune response, T cell activation, and stimulus response indicate a shared TFH program is imprinted through antigen-experience and successive B cell contacts. Largely limited to the post-GC day 28 TFH, we find catabolic metabolism, apoptosis prevention, and transcription reduction (**Figure 6 C**). We additionally see GO enrichment of regulatory cytokine (*TFGβ, Il-10 and IFN-γ*) production (**Figure 6 C**). Taken together, the day 28 TFH transcriptional profile indicates the post-GC compartment is surviving in a decreased transcriptional and metabolic state and may serve limited regulatory functions.

While we see a conservation of, and sort on, CXCR5 expression, we see a significant loss of PD-1 surface expression (**Figure 6 D, E**). We found that 40% of sorted day 28 TFH had lost PD-1 expression, and as a key TFH identifier thought to be most highly expressed in the GC, we further investigated the transcriptional profile to uncover systematic shifts linked to PD-1 downregulation (**Figure 6 F**). Unsurprisingly, the PD-1 high population is similar to the primary day 14 TFH with the shared expression of transcription factors (*Bcl6, Pou2af1* (Bob1), *Cited2, Maf, Ascl2*) and surface molecules (*H2-K1, Tnfsf11, Sh2d1a, Entpd1, Cd40lg*) (**Figure 6G**, **S6 D**). Despite these common features, regulatory (*Il10, Vdr, Havcr1*, *Smad2*) and cell-cycle arrest (*Plac8, Crebbp, Tle1, Lef1*) associated genes appear upregulated in the PD-1 high population (**Figure 6 F, G**). Taken together, the PD-1 high day 28 TFH appear to be a non-proliferative regulatory phase of TFH function with reduced cellular metabolism and transcription.

In the PD-1 low compartment the expression of regulatory associated genes (*Il10, Vdr, Havcr1, Smad2*) and transcription factors (*Bcl6, Pou2af1, Ascl2*) is lost suggesting the quiescence of the Bcl6 driven germinal center program (**Figure 6 F, G**, **S6 E**). However, there is continued expression of *Tle1, Ltbp3,* and *Cd28* in addition to the upregulation of an alternative inhibitory pathway (*Sigirr, Tgfb3, Timd2, Lag3, Btg1*). Further distinguishing the PD-1 low population are elevated levels of cell cycle (*Gfi1, Ccnd2, Jag1*), metabolism (*Slc2a1, Mxd1, Gclc*), signaling (*Tlr7, Tnfsf14, Eef2, Nfat5*), and adhesion (*Lsp1, S1pr1*) associated genes. These features suggest PD-1 low TFH are programmed for continued persistence and primed for secondary antigen exposure.

Supporting a stepwise progression from primary day 14 TFH, through a regulatory PD-1 high day 28 TFH, to a PD-1 low memory day 28 TFH, we find the PD-1 low population shares transcriptional features with both TCM and TEM populations. The upregulation of TCM associated genes (*Thy1, Foxj1*) exclusively in the PD-1 low population indicate a potential link between local memory TFH and circulating CXCR5+ TCM (**Figure 6F**, **S6 F**). Conversely, the PD-1 low population also upregulates genes expressed in the TEM cohort (*Irf4, Prkdc, Nfkbia*) (**Figure 6 F**, **S6 F**) potentially indicating the existence of follicular memory within both TCM and TEM compartments. In culmination, we consider the PD-1 high and low post-GC transcriptional programs characteristic of a progressive evolution of TFH function from primary activity to the conclusion of effector function before finally entering a long-lived memory state.

## Discussion

The management of the antigen-specific B cell response is essential for the protection and maintenance of health. Our studies of antigen-specific TFH, responsible for orchestrating high-affinity B cell immunity, demonstrate the clonal plasticity of naïve CD4 T cells and the significant influence of affinity on TFH development. We propose inflammatory and anti-inflammatory transcriptional patterns, complementary to a core follicular program, distinctly specializing TFH to provide class-specific B cell support. Furthermore, we show this follicular program passes through a regulatory stage prior to transitioning into a quiescent local memory state while maintaining core TFH features.

The extent of naïve CD4 T cell plasticity and the influence of TCR affinity has been a long-standing debate (*43, 46-48*). Higher affinity for antigen results in more cognate contact, both in duration and frequency (*4, 49, 50*). Monoclonal transgenic studies have shown robust expansion into multiple distinct T cell lineages (*48, 51*), however these systems, by design, lack clonal diversity and are thus better suited to investigate the role of affinity rather than plasticity. Recent work sequencing over 600 individual LCMV specific clones showed the majority of clones differentiate into both T_H_1 and TFH; however, roughly 30% were exclusively T_H_1 or TFH (*46*). Among the exclusive clones are favored motifs and these preferred structural characteristics may influence affinity (*46*). Our data similarly show the expansion of clonotypes in both conventional effector T_H_ (where T_H_1 reside) and TFH populations, and through Jβ usage, found a distinct skewing of fate linked to TCR affinity. This aligns with previous work illustrating TFH differentiation is correlated with increased TCR affinity, signaling duration and density, and frequency of contacts (*43, 47, 52*). However, recent work illustrates high-affinity clones favoring T_H_1 differentiation during acute LCMV infection and conversely favoring the TFH fate during chronic infection (*48*). This is among work which suggests the context of activation influences the role of affinity in fate determination (*48, 51*).

We instead see the influence of activation context in the shared inflammatory transcriptional program across ETH and TFH. The consistent upregulation of inflammatory gene programs reflects the type 1 skewed activation of dendritic cells driven by the MPL adjuvant, a TLR4 agonist (*53*). While others have described a transitional co-expression of Tbet (T_H_1) and Bcl6 (TFH) (*22, 23*), we find a sustained inflammatory panel and downstream inflammatory B cell products demonstrating a cascade of effector function communicated through sequential bi-directional programming.

Where previous work has shown either type 1 or type 2 skewing TFH profiles, these studies were in singularly type 1 or type 2 models without concurrent representation of both the inflammatory and anti-inflammatory response (*22-25, 29, 54*). We show the specific and simultaneous presence of pro-inflammatory and anti-inflammatory transcriptional signatures within the TFH compartment throughout the primary B cell response; directly correlating with inflammatory IgG2b and anti-inflammatory IgG1 B cell maturation. This demonstrates that inflammatory and anti-inflammatory subspecialization of TFH can co-exist and occur in the primary immune response, not just under conditions of chronic and intense polarization (*24, 25, 54*). As our recent work demonstrates the capacity for a single surface ligand (PD-1) to have extensive and exclusive class-specific (IgG1) impact (*44*), these transcriptional profiles provide a robust array of targets for CD4 mediated modulation of class-specific responses.

With the collapse of the primary antigen-specific GC, we uncovered the transcriptional progression from primary effector function into memory. The consistent expression of inhibitory receptors (*Pdcd1, Havcr1, Rgs10, Tnfsf8, Tigit, Ctla4*) suggests the evolutionary importance of tightly regulating B cell maturation and reflects the ability to quickly shut down TFH activity to avoid autoimmune dysregulation via hyperactive GCs (*55*). We found that the local post-GC TFH exhibit robust inhibitory transcriptional patterns distinct from TEM and TCM. Interestingly, recent work indicated GC TFH upregulate regulatory transcription factor, *Foxp3*, to mediate the collapse of the GC (*56*). While we did not find differential expression of *Foxp3*, four of the most significant day 28 TFH distinct genes – *Skil, Smad3, Ltbp3,* and *Vdr*– are associated with regulating TGFβ signaling: TGFβ1 has been shown to induce *Foxp3* expression(*57*). TGFβ1 is also instrumental in maintaining T cell homeostasis; preventing apoptosis, limiting proliferation, dampening cytokine production, and avoiding autoimmunity (*58*).

We see the loss of this primary inhibitory pattern and *Bcl6,* in addition to the upregulation of metabolism, cell-cycle, and survival associated genes in the PD-1 low day 28 TFH indicating non-GC function and the capacity for long-term persistence. The PD-1 low TFH population appears more programmatically similar to the TCM and TEM populations than the primary day 14 TFH and thus we propose this population is antigen-specific memory TFH. We see the upregulation of TCM markers (*Thy1, Foxj1*) reflecting the phenotype of circulating memory CXCR5+ T cells (*21, 34*); however, studies of CXCR5+ TCM also see maintained expression of PD-1 (*33*). Conversely, we also see the shared expression of metabolic factors between PD-1 low TFH and TEM cohorts. The similarities to both TCM and TEM suggests a possible divergence of follicular memory into both circulating and resident memory fates.

This work indicates the TFH population is as dynamic and heterogeneous as the B cell population they regulate. We found robust inflammatory and anti-inflammatory modules of transcriptional function correlated with inflammatory and anti-inflammatory B cell maturation. While TFH subsets have been discussed in the field, we provide clear evidence that specialized TFH function is present in healthy immune responses, not only dysfunctional or chronic immune challenges. These arrays of type 1 and type 2 associated molecules expressed on antigen-specific TFH will inform future vaccine design to direct class-specific support and therapies targeting class-specific dysregulation. Furthermore, these connections may inform potential off-target humoral effects as many lymphoid molecules are promising drug targets.

Lastly, we illustrate the stepwise progression of TFH from primary GC function through GC collapse and into memory. We propose the transitional expression of surface ligands and signaling molecules may aid in the identification and study of memory TFH prior to restimulation upon secondary exposure. The survival and maintenance of the memory TFH population is crucial to confer long-term protection against challenge. Understanding the transcriptional programming of long-lived TFH memory will inform vaccine strategies and CD4 based CAR-T therapies. These findings underscore the functional heterogeneity within the TFH population; it will be important to extend these pattern-based findings into disease models and therapeutic development.

## Materials and Methods

### Mice

B10.Br mice were bred and housed in specific pathogen-free conditions. All experiments were done in compliance with federal laws and institutional guidelines as approved by The Scripps Research Institutional Animal Care and Use Committee.

### Immunizations

Mice were given primary immunization subcutaneously at the base of the tail with 400μg whole PCC (Sigma) or 400µg NP-PCC (4-hydroxy-3-nitrophenylacetyl (Biosearch) conjugated to pigeon cytochrome C (Sigma)) mixed with adjuvant based on Monophosphoryl Lipid A, lab formulated based on ref (*59*). MPL-based adjuvant was made in house by emulsifying Monophosphoryl Lipid A [Synthetic] (Avanti) with lecithin (Spectrum) and Tween-80 (Sigma-Aldrich) in squalene (Sigma-Aldrich) based on the formulation described by Baldrige and Crane (*60*). NP-PCC was conjugated in house using NP-Osu (Biosearch) and Pigeon Cytochrome C (Sigma). Mice were selected for analysis on the basis of days elapsed beyond the initial date of immunization, defined as day 0.

### Flow cytometry

Draining lymph nodes (inguinal and periaortic LNs) were removed from unimmunized (Naïve) or immunized (PCC or NP-PCC in MPL) mice and passed through 70μM cell strainers (Falcon) in ACK lysis buffer (Gibco) while on ice. Single-cell suspensions were then prepared in “FACS wash” made in-house from Phenol red-free DMEM (Gibco) and 2% (vol/vol) FBS. Cells (4 × 10^8^ per ml) were incubated on ice for 10 min with anti-CD16/32 ‘Fc block’ (2.4G2; Bio X Cell) prior to staining in the dark for 30 min on ice with fluorophore or biotin-labeled monoclonal antibodies prepared in brilliant stain buffer (BD Biosciences) ‘cocktail’ at pre-titrated quantities. Streptavidin-conjugated secondary antibodies were used as a second-step visualization reagent for biotin-labeled antibodies, see Table S1 for monoclonal antibodies used.

For surface staining of IgG1 and IgG2b non-specific Fc binding was blocked with anti-CD16/32 (identified above) for 10 minutes on ice. Before addition of secondary, antibody cocktail, non-specific immunoglobulin binding was blocked by addition of 2% (vol/vol) normal mouse and normal rat serum (1:1 mix) in FACs Wash. Pre-titrated immunoglobulin specific antibody cocktails prepared in brilliant stain buffer (BD) were then added for 30 min on ice. Cells were washed with FACS wash and resuspended in 2% normal mouse and normal rat serum (1:1 mix) before cells were stained with a ‘cocktail’ containing fluorophore-conjugated antibodies for 30 min on ice.

High-resolution analysis with a four-laser FACS Aria III with index cell sorting was central to all studies. The use of 12- to 15-color resolution for analysis and isolation of single antigen-specific T and B cells provided the requisite purity, as attested by efficiency of single-cell gene expression. Analysis was done on a FACS Aria III with FACS Diva 8.0 software equipped with index sorting software (BD Biosciences) for recording of the exact data for each single cell sorted into each well of a 96 or 384-well plate. Indexed cell sorting permitted the direct tracking of individual cells processed for surface phenotype and gene expression to quantify the levels and penetrance frequency of functionally interesting genes. Flow cytometry data were analyzed with FlowJo software (TreeStar).

### Single-cell T cell receptor (TCR) repertoire analysis

Individual lymph nodes day 7 immunized mice were collected and single cells with the appropriate surface phenotype sorted by flow cytometry into 96-well plate plates containing 5μl RT-Pre-Amplification Master Mix (2.5μl CellsDirect 2X Reaction Mix; 0.1μl SuperScript III RT/Platinum Taq (CellsDirect One-Step qRT-PCR kit; Invitrogen); 0.25μl pooled 0.5μM external Vb3 primers (Vb3; 5’ ATG GCT ACA AGG CTC CTC TGG TA 3’ and 5’ CAC GTG GTC AGG GAA GAA 3’); 2.15μl diethylpyrocarbonate-treated double-distilled water) (as previously described (*30*)). As negative control for each plate, at least four wells per plate had no cell and were processed throughout the procedure. The reverse transcription was performed at 50 °C for 15 min, followed by 95 °C for 2min, then 22 cycles of 95 °C for 15s, 60 °C for 4min. The amplified cDNA was diluted five times in Tris-EDTA buffer, then 1μl of diluted product used for nested PCR in a volume of 10μl containing 2U/ml Taq polymerase, 200μM of each dNTP (Roche), 1mM Tris-HCl, 1.5mM MgCl, 25mM KCl and 0.8μM of internal primers (Vb3; 5’ AAT CTG CAG AAT TCA AAA GTC ATT CAG 3’ and 5’ AAT CTG CAG CAC GAG GGT AGC CTT TTG 3’) for amplification of the gene encoding the Vb3 with the PCR program: 95 °C for 5 min then 40 cycles of 95 °C for 15 s, 55 °C for 45s, 72 °C for 90s, ending with 72 °C for 5 min. PCR products were purified (ExoSAP-IT; USB) and then were sequenced with a Big Dye Terminator v3.1 cycle sequencing kit (Applied Biosystems) with 3.2pmol of the second-round PCR primer for Vb3 (5’ CTG TGC TGA GTG TCC TTC AAA C 3’) products. Sequence products were purified (BigDye XTerminator purification kit, Applied Biosystems) and run on a 3130 genetic analyzer (Applied Biosystems). Gene segments were assigned as encoding variable, diversity and joining regions through the BLASTn tool (National Center for Biotechnology Information).

### Microdissection

Draining lymph nodes (inguinal and periaortic LNs) removed from unimmunized (Naïve) or immunized (NP-PCC in MPL) mice were kept on ice in ACK lysing buffer (Gibco). Under a dissecting microscope (Nikon, C-BD115 + high intensity illuminator NI-150) nonlymphoid tissue was removed from the surface with forceps prior to dissection. Each lymph node was divided into two with a clean razor blade and processed independently as described above. Staining volumes were adjusted to the cell counts (4 × 10^8^ per ml). The entirety of each LN portion was stained and run through FACS.

### ELISpot

As described previously (*18*), 96 well PVDF-bottomed plates (Millipore #MAHA-S4510) were washed with filtered PBS prior to coating with NP23 -BSA (conjugated in-house, 25μg/mL) or goat anti-mouse IgM/IgG/IgA antibody heavy and light chain (25μg/mL); IgM (AF6-78; Biolegend), IgG1 (RMG1-1; Biolegend), IgG2b (RMG2b-1; Biolegend), IgG2a (R19-15; BD bioscience), IgG3 (RMG3-1; Biolegend), IgA (RMA-1; Biolegend). Wells were blocked by incubating DMEM media (Gibco) containing 5% FBS for 1 hour at 37°C. Bulk unstained cells from the dLN suspended at 4x106 cells/mL in DMEM (Gibco) with FBS (5% vol/vol) were added to wells at serially titrated quantities (10x104 to 1.25x104 cells). Negative control wells were included, for which medium only was plated. Plates were incubated at 37°C for a minimum of 14 hours, then wells were then washed prior to addition of horse radish peroxidase (HRP) conjugated goat anti-mouse Fc-specific antibodies for 4 hours at room temperature. [1:500 dilution for IgM and IgG subclasses, 1:300 for IgA.] HRP conjugated antibodies were all from Southern BioTech: IgM (#1021-05), IgG (#1030-05), IgG1 (#1071-05), IgG2b (#1090-05), IgG2a (#1081-05), IgG3 (#1100-05), IgA (#1040-05). Wells were washed and spots were detected by adding a filtered solution of 0.2mg/mL AEC (3-amino-9-ethylcarbazole; Sigma) in 0.1M sodium acetate buffer [pH 5.0] and 0.05% hydrogen peroxide (Sigma). Wells were washed and air dried for a minimum of 24 hours prior to counting the spots by visual inspection under a stereomicroscope.

### Single cell Quantitative and Targeted single cell RNAseq (single cell qtSEQ)

The following method is detailed in ref. (*45*) and in practice in ref. (*44*). Single cells were index sorted into 384 well plates containing 1uL of reverse transcription (RT) master mix using a FACS Aria III with FACS Diva software (BD Biosciences) to maintain protein expression with plate well location. The RT master mix consisted of: 0.2μL SuperScript IV 5x buffer (Invitrogen), 0.03μL SuperScript IV taq (Invitrogen), 0.03 DTT 0.1M (Invitrogen), 0.03μL RNaseOUT (Invitrogen), 0.012μL dNTPs (Invitrogen), 0.19μL RNase Free H2O and 0.5μL of oligoDT primer [1μM stock] (custom made, IDT) per well. A unique oligoDT primer that contains a 6 basepair well barcode and a 10 54 basepair unique molecular identifier was added to each well. Reverse transcription was performed at 42°C for 50 minutes followed by heat inactivation at 80°C for 10 minutes.

All 384 wells were pooled following RT, for the removal of excess oligoDT primers using ExoSAP-IT express (Affymetrix). After pooling, 0.3μL of Exo-SAP-IT express was added per 1μL of pooled library; thermocycling conditions were 20 minutes at 37°C followed by heat inactivation at 80°C for 2 minutes. Volume reduction and additional oligoDT removal was performed using AMpureXP SPRIselect paramagnetic beads (BeckmanCoulter) at a 0.8:1 bead to library ratio with our pooled library. To reduce the quantity of paramagnetic beads used per plate, 100uL of beads were used per library with pure bead binding buffer for the remaining volume (Bead Binding Buffer, 2.5M NaCl, 20% vol/vol PEG, Teknova #P4146). The beads and library were mixed and allowed to bind for 5 minutes at room temperature before using a magnetic stand to pellet the beads for removal of supernatant. A wash of 200μL 85% (vol/vol) molecular grade ethanol was added for 30 seconds, then removed and beads were allowed to air dry for ∼2 minutes. The cDNA was eluted off the beads by 35uL of PCR-grade water.

A nested PCR approach was used for targeted gene amplification. RNaseH treatment to remove mRNA from the cDNA preceded PCR targeted gene mRNA amplification. PCR1 utilized a singular reverse primer dubbed “RA5-overhang” (5’ AAG CAG TGG TGA GTT CTA CAG TCC GAC GATC 3’) and selected gene specific primers that targeted the 3’ terminus in gene coding sequence of the untranslated region (UTR). PCR1 was performed in a 60μL reaction: 35μL cDNA library, 1.5μL RNaseH (NEB), 1.2μL Phusion DNA polymerase (NEB), 12μL 5X Phusion HF Buffer (NEB), 0.85μL dNTPS [stock 25μM] (Invitrogen), 1.2μL of RA5-overhang [stock 100μM], and 25nM final concentration for each specific gene targeting external primer. The volume of gene specific primers varied depending on number of specific primers used, and water was added to reach a 60μL reaction volume if needed. PCR1 template amplification thermocycling was as such: 37°C for 20 minutes followed by 98°C for 3 minutes for RNaseH activity and heat inactivation, followed by 10 cycles of 95°C for 30 seconds, 60°C for 3 minutes, 72 °C for 1 minute. Completion of ends was performed via 72°C for 5 minutes. Removal of excess primers and volume reduction was performed again using AMpureXP SPRIselect beads, at a 0.9:1 bead to library ratio, and eluted in 20μL of water following the same protocol as listed above.

PCR2 utilized nested primers binding sites internal to PCR1 primers. A singular reverse primer was used, RA5 [5’ GAGTTCTACAGTCCGACGATC 3’] and selected gene specific primers that targeted the 3’ terminus in gene coding sequence of UTR. These “Internal” primers also contained an overhang sequence for Illumina flow cell binding adapters to be added. [See ref (*45*)]. Half of the PCR1 library elute was used in PCR2 template amplification such that PCR2 contained 10μL PCR1 reaction elute, 0.4μL Phusion DNA polymerase (NEB), 4μL 5X Phusion Buffer (NEB), 0.3μL dNTPs [stock 25μM] (Invitrogen), 2μL of RA5 [stock 20μM], and 25nM final concentration for each specific gene targeting internal primer. The volume of gene specific primers varied depending on number of specific primers used, and water was added to reach 20μL reaction volume if needed. PCR2 conditions were as such: 95°C for 3 minutes followed by 10 cycles of 95°C for 30 seconds, 60°C for 3 minutes, 72 °C for 1 minute. Completion of ends was performed via 72°C for 5 minutes. Removal of excess primers and volume reduction was performed again using AMpureXP SPRIselect beads, at a 0.9:1 bead to library ratio and eluted in 20μL of water following the same protocol as listed above.

PCR3 was performed to add library identifying sequences necessary for sequencing multiplexing as well as Illumina flow cell binding adapters. For each library, a unique plate identifier primer (RPI primers) was added with a shared primer RA1. Half of the PCR2 library elute was used in PCR3 template amplification such that PCR3 contained 10μL PCR2 reaction elute, 0.4μL Phusion DNA polymerase (NEB), 4μL 5X Phusion Buffer (NEB), 0.3μL dNTPs [stock 25μM] (Invitrogen), 2μL of RP1 [stock 10μM], 2μL of RPI primer [stock 10μM], and 1.3μL water. Half of the PCR2 library elute (10μL) was set aside and saved. PCR3 template amplification thermocycling was as such: 95°C for 3 minutes followed by 8 cycles of 95°C for 15 seconds, 60°C for 30 seconds, 72 °C for 30 seconds. Completion of ends was performed via 72°C for 5 minutes. Removal of excess primers and sample purification was performed using AMpureXP SPRIselect beads at a 0.7:1 bead to library ratio and eluted in 20μL of water following the same protocol as listed above.

The prepared cDNA library concentration was quantified using the Qubit dsDNA HS kit (Invitrogen) and library peak sizes determined via a high sensitivity DNA chip run on a bioanalyzer (Agilent). Conversion of qubit concentration (ng/μL) to molarity was performed using the prescribed calculation from Illumina.

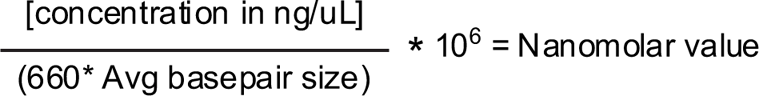

Preparation of libraries for sequencing followed the prescribed steps outlined in the Illumina Denature and Dilute Libraries Guide (Illumina). In short, DNA libraries were denatured using 0.2M NaOH, quenched with 200mM Tris-HCl, and diluted to 1.8pM in a final volume of 1.3mL solution of provided HT1 buffer for loading onto a sequencing chip. Libraries were sequenced on a NextSeq500 (Illumina) using V2 sequencing chips (Illumina). A paired-end read setting (75bp) was used to cover barcode and UMI, plate identifier sequence (RPI) and sequencing of 3’ UTRs. For sequencing, the following numbers of cycles were used: Read 1: 15 cycles, Index Read 1: 6 cycles, Read 2: 71 cycles.

### Data processing, quality control, and analysis

Bcl2fastq (Illumina) was used to demultiplex sequencing libraries using plate barcodes and assort reads by well barcodes prior to sequence alignment using Bowtie2 (v2.2.9) to a custom genome consisting of the amplicon regions for the specifically targeted genes used in the experiment based on the murine genome GRCm38.p6, version R97 (Ensembl). HTseq was used for read and UMI reduced tabulation. Custom scripts for these steps available upon request.

Quality control excluded cells that contained over 2000 UMIs and fewer than 1000 reads, 60 UMI counts, and 25 unique genes detected. A secondary quality control filter included T cells that contained expressed Cd3e, Cd3d, or Cd3g. A total of 1924 T cells from 5 experiments were analyzed.

Data from the two separate experiments was pooled into a single count matrix for processing using the R package Seurat (v3.2.3) (*61*); scaling and normalization was carried out using the SCTransform package (v0.3.2.9002, (*62*)) with the batch_var option set to the metadata variable “Experiment”. This normalized UMI count matrix was then used for downstream analysis. SCTransform was additionally executed on naïve, ETH and TFH cell types separately for day 7 analysis (Fig2).

For differential gene expression analyses, MAST (Model-based Analysis of Single Cell Transcriptomics) package (v1.16.0 https://github.com/RGLab/MAST/ (*63*)) using a hurdle model was used to generated fold change and p-values for all genes. Volcano plots were then created using Prism v9.0 (GraphPad Software). The R package pheatmap (v1.0.12) was used to plot heatmaps of averaged UMI counts. Single cell heatmaps were generated using the “Doheatmap” function from the Seurat package. Uniform Manifold Approximation and Projection (UMAP) was performed using “RunUMAP” function from the Seurat package. UMAP function options; “dims” was set to the top 3 principal components following PCA and “metric” set to euclidean distance. Custom scripts for these steps available upon request.

### Statistical analysis

Statistical analysis performed in R detailed above. All additional statistical analysis was performed using GraphPad Prism v 9.0. Statistical tests and number of replicates are reported for each experiment.

### Data and Materials Availability

All customized R codes used for data analysis and raw and processed data files for the scRNAseq analysis in the study will be provided upon request without restriction.

## Acknowledgements

This work was supported by the US National Institutes of Health (AI047231, AI040215 and AI071182) and Bill & Melinda Gates Foundation (BMGF 0PP1154835) to M.G.M.-W.

## Author contributions

A.M.R, B.W.H, A.G.S, K.B.M, L.J.M-W, and M.G.M-W designed and performed experiments, B.W.H, A.G.S, L.J.M-W, and M.G.M-W designed and developed single cell qtSEQ, A.M.R, L.J.M-W, and M.G.M-W designed the study, analyzed data and wrote the paper.

## Supplementary materials

**Fig S1.**
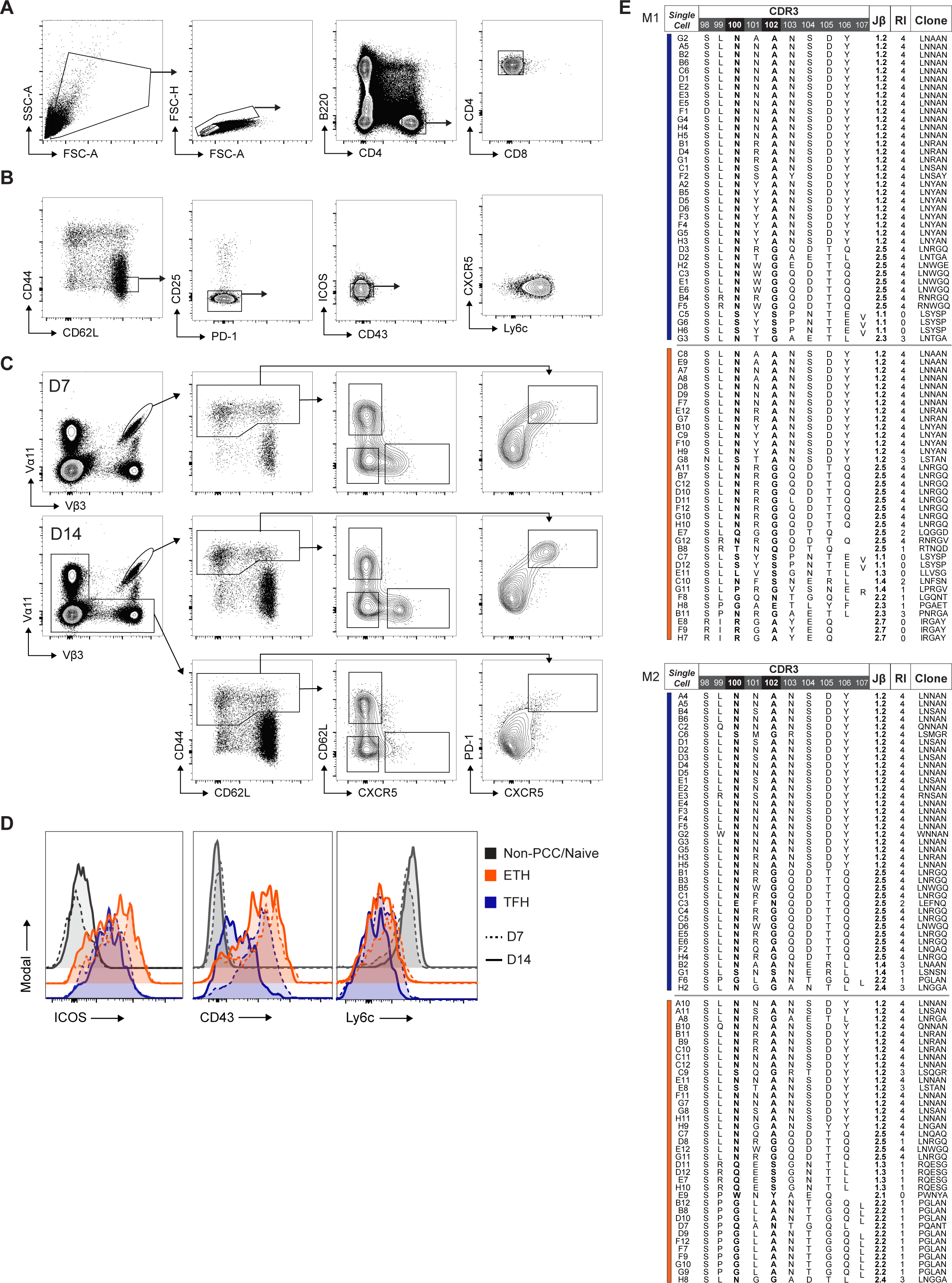
Extended intraclonal analysis of PCC specific TFH and ETH. **(A)** FACS gating strategy to isolate CD4+ T helper cells. **(B)** FACS gating strategy for non-PCC specific “Naive” population. **(C)** FACS gating strategy for activated CD4+ populations, Va11Vb3+ PCC specific (D7 and D14) and non-specific (representative from D14 mouse). **(D)** FACS histogram of ICOS, CD43, and Ly6c expression. Dashed lines represent D7 sample; Solid lines represent D14 sample. Black represents non-PCC MPL background “naïve” CD62L+ CD44-TH, Orange - Emmigrant T_H_ (ETH), Blue - TFH. **(E)** Vβ3 CDR3 region sequences of index sorted Vα11 Vβ3 CD44hi of other 2 mice matching Fig 1 D.

**Fig S2.**
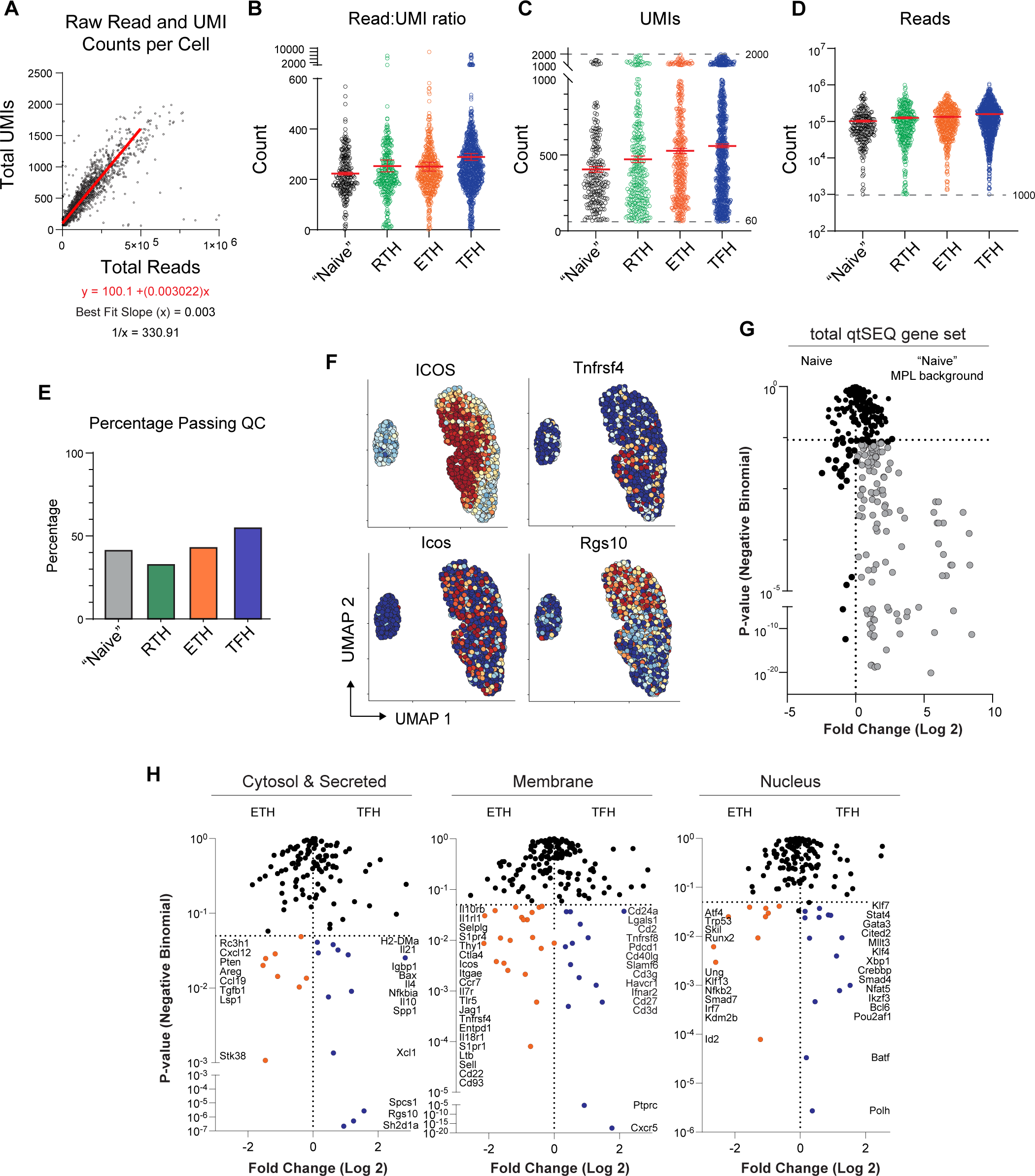
qtSEQ biological and technical requirements. **(A)** Reads vs UMI counts of each sequenced cell that passed QC requirements. Best fit first-degree polynomial line; 500<x<500000. **(B)** Read:UMI ratio grouped by index sorted phenotype. **(C)** UMI count grouped by index phenotype. Dashed lines indicate technical thresholds. **(D)** Read count grouped by index sorted phenotype. Dashed line indicates technical threshold. **(E)** Percentage of indicated cell populations which passed biological and technical QC thresholds. **(F)** Transcriptional and protein expression overlayed on UMAP dimensionality reduction of index sorted “Naïve” and antigen-specific ETH and TFH. **(G)** Differential gene expression between Naive (unimmunized spleen) and “Naive” MPL background. **(H)** Differential gene expression between ETH and TFH, populations separated by designated location/function. Fold change plotted against negative binomial P-value

**Fig S3.**
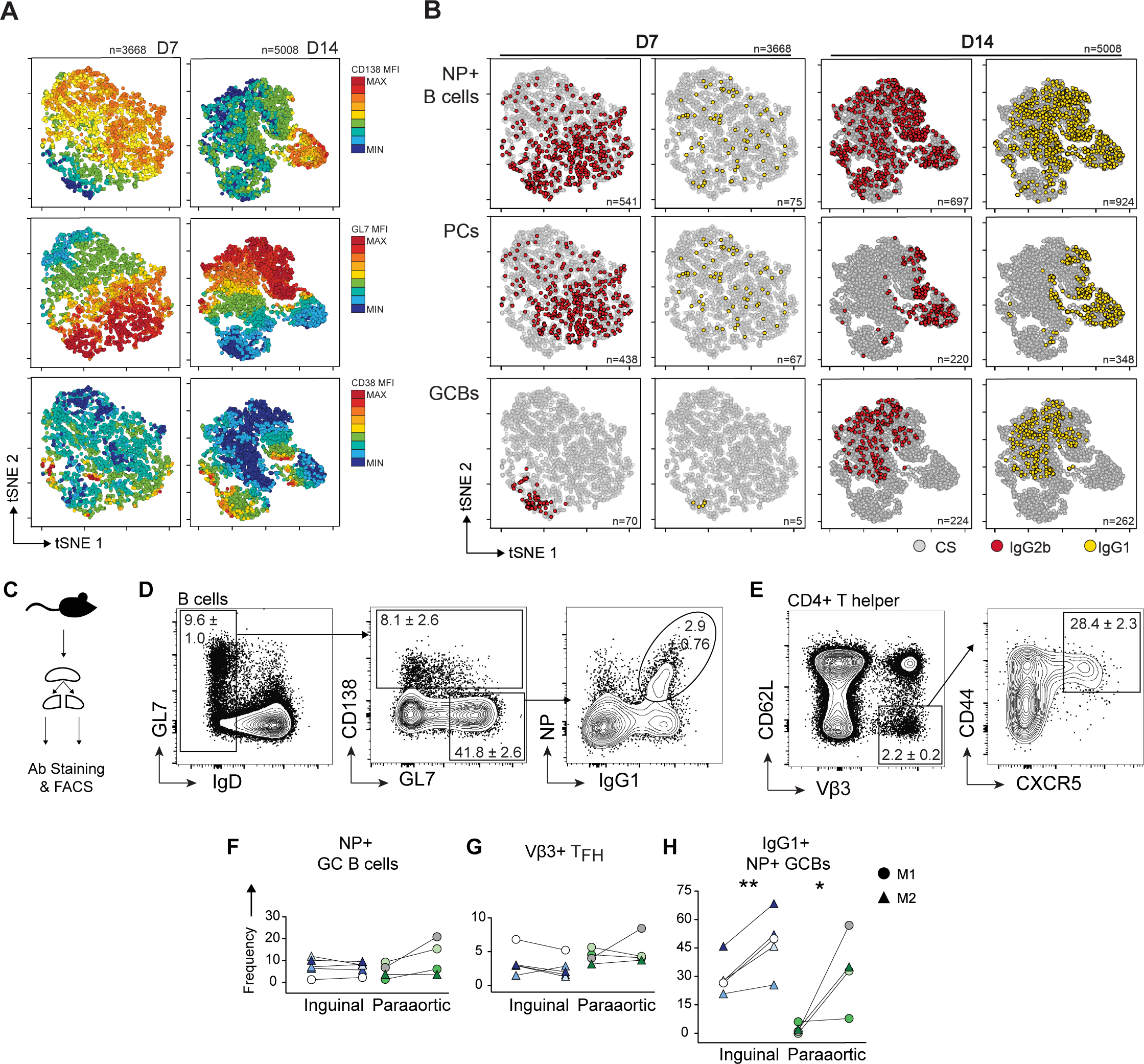
Temporally delayed and geographically skewed anti-inflammatory B cell response. **(A)** CS antigen-specific B cell (CD3-, Gr1-, CD19 and/or CD138+, IgD-, IgM-, NP+, L1+) D7 and D14 tSNE map protein expression of CD138, CD38, and GL7. **(B)** IgG2b and IgG1 expression overlayed on tSNE at D7 and D14. Cell count of IgG2b and IgG1 B cells indicated in lower right corner of plot. Total plotted cells indicated above. **(C)** Microdissection and FACS strategy schematic. **(D)** Representative flow plots show the isolation of antigen specific IgG1+ GC B cells 14 days post NP-PCC + MPL immunization. IgD-CD19+ B cells are separated as CD138 + PCs and GL7+ GCB cells. NP-specific IgG1 GCB cells are isolated from the GL7+ GC population. **(E)** Gating strategy isolating Vβ3+ TFH within the same stain shown isolating NP-specific B cells in (D). **(F-H)** NP-specific GC B cell **(F)**, Vβ3+ TFH **(G)**, and NP-specific IgG1 **(H)** frequencies within separate regions of the same inguinal or periaortic LN. Data from 2 mice: 3 inguinal, 4 periaortic LNs. Data points of the same color are from the same LN section. Paired one-tailed t.test, * P≤0.05, ** P≤0.01.

**Fig S4.**
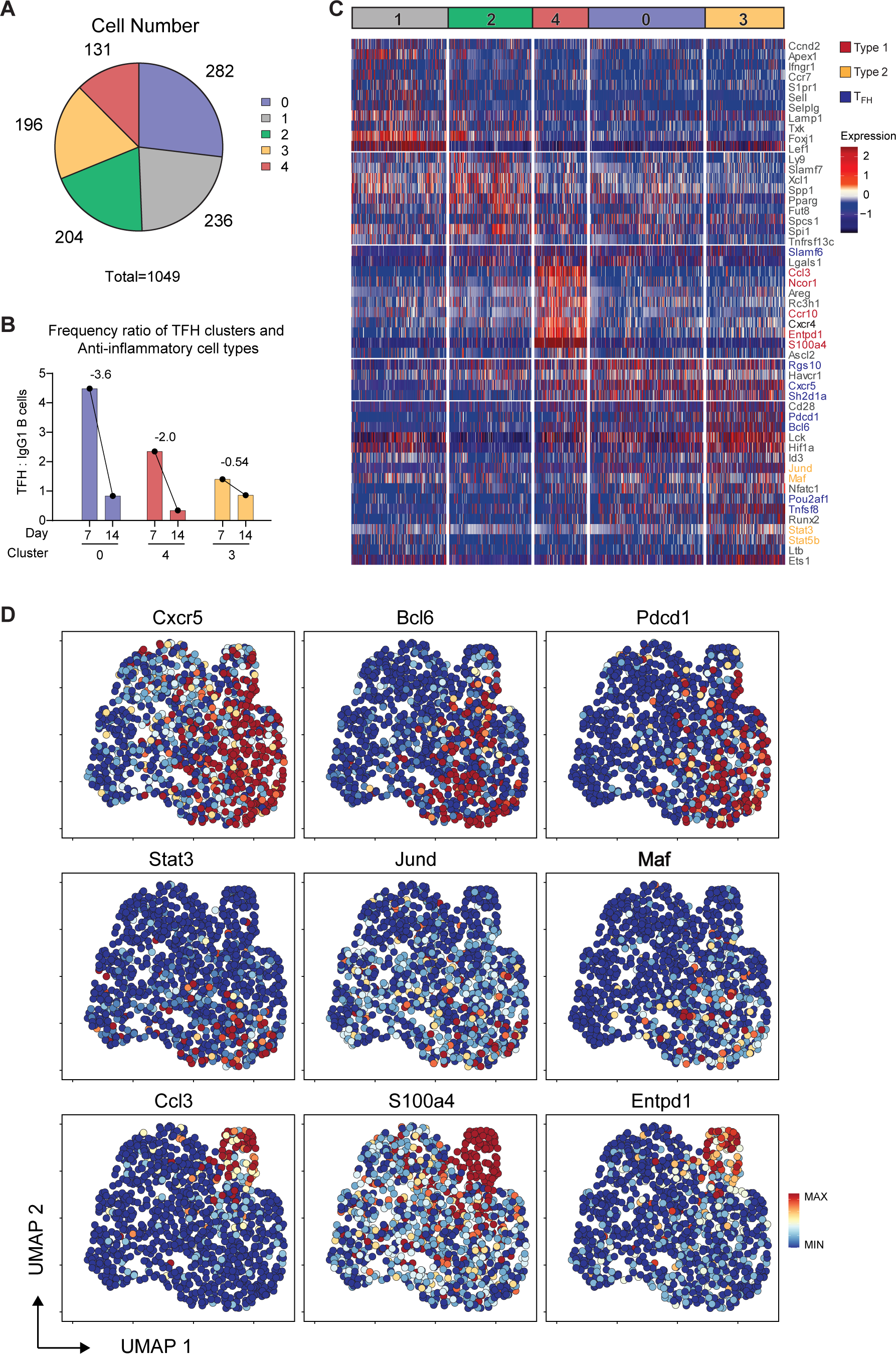
Extended Inflammatory and Anti-inflammatory transcriptional analysis. **(A)** Cluster proportions of total sample. Cell count of each cluster listed. **B)** Single cell gene expression heat-map. **(C)** Frequency ratio of TFH clusters and anti-inflammatory cell types at D7 and D14, bar at median of B cell measurements. Relative slope between D7 and D14 ratios listed. **(D)** Heatmap expression of differentially expressed genes on UMAP coordinates. Genes^55^are as labeled.

**Fig S5.**
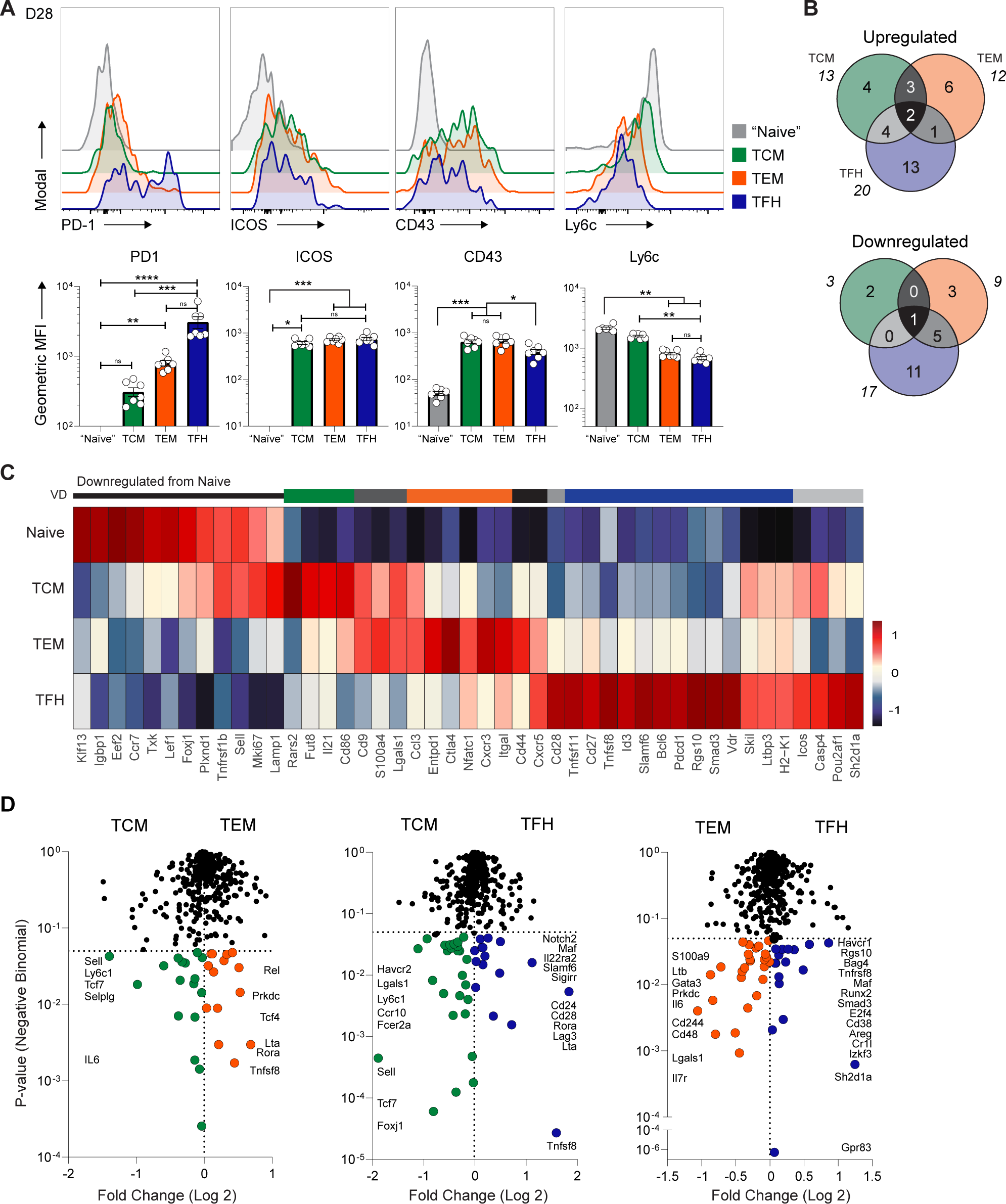
Extended putative local memory TFH analysis. **(A)** Representative FACS histogram of PD1, ICOS, CD43, and Ly6c expression and MFI quantification of respective surface markers. Black represents Naive TH, Blue - Ag-specific TFH. (Uncorrected Dunn’s test, P<0.05 *, P<0.01 **). **(B)** Venn Diagram of shared and unique gene expression across TCM, TEM, and TFH populations. Genes have P-value <0.05, and log2 fold change >0.1. **(C)** Pseudobulk heatmap expression of genes represented in venn-diagrams (B) across Naive/Non-PCC, TCM, TEM, and D28 TFH populations. Colored side bar (VD) represents the region of the venn-diagram represented. **(D)** Differential gene expression between TCM, TEM, and D28 TFH populations. Fold change plotted against negative binomial P-value.

**Fig S6.**
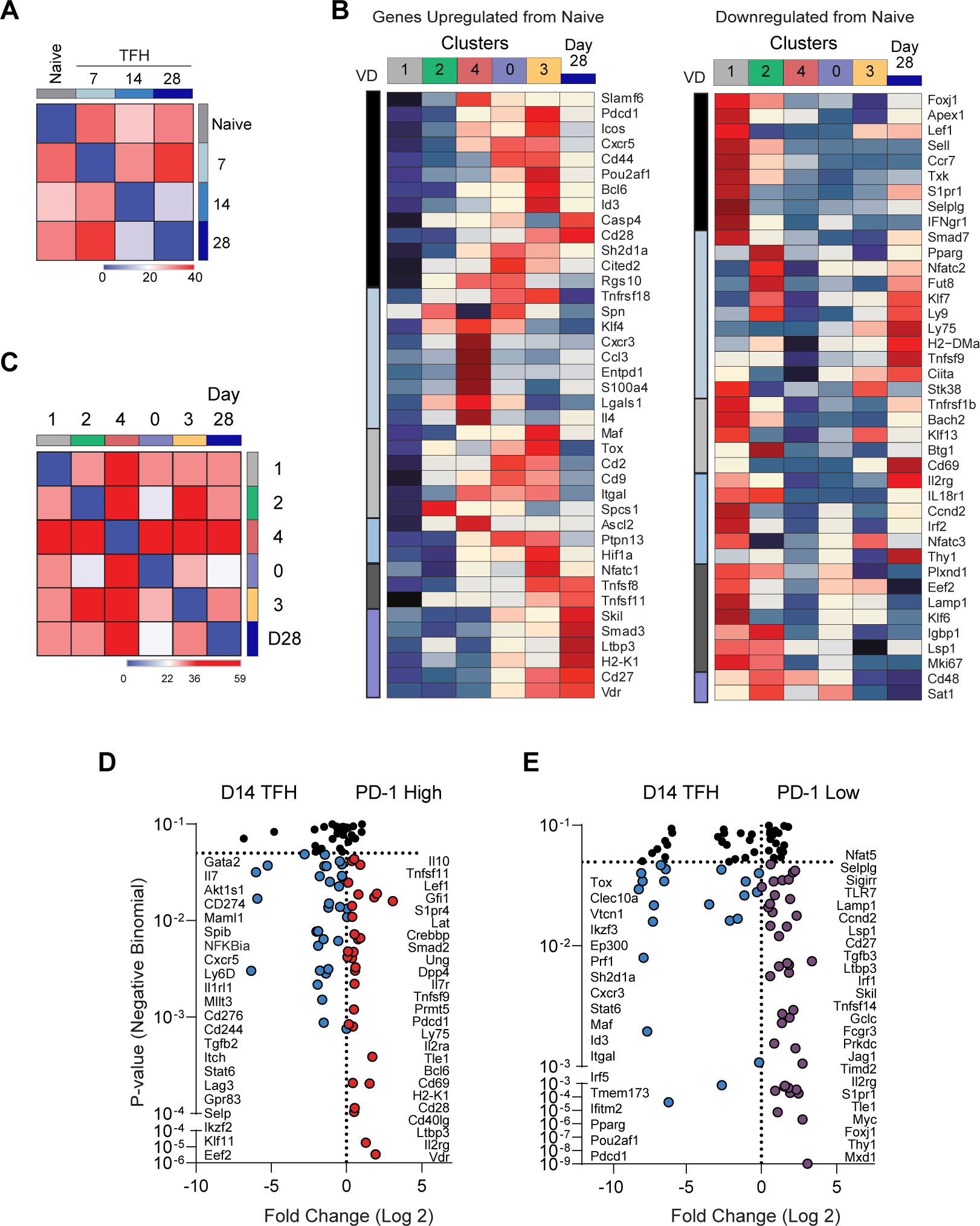
Extended memory transcriptional analysis. **(A)** Euclidean distance similarity matrix across Naive TH, D7, D14, and D28 TFH. **(B)** Pseudobulk heat-map of genes listed in up- and downregulated venn-diagrams. VD column corresponds to the area of the venn-diagram represented. Heatmap of Figure 4 UMAP clusters and D28 PCC-TFH. (**C)** Euclidean distance similarity matrix across UMAP clusters and D28 TFH. **(D)** Differential gene expression between primary D14 TFH and PD1 high (D) or PD1 low (E) D28 TFH populations. Fold change plotted against negative binomial P-value.

